# From Coral Color to Tissue Heating: How Host Pigments Redirect Light Within the Tissue

**DOI:** 10.64898/2026.10.20.760344

**Authors:** Swathi Murthy, Elena Bollati, Anya Salih, Michael Kühl

## Abstract

Coral host pigments strongly modify the internal light environment of corals, yet their influence on heat dissipation in coral tissue remains poorly understood. Here, we combined scalar irradiance and temperature microsensor measurements with mechanistic multiphysics modelling to quantify how chromoproteins (CPs) and fluorescent proteins (FPs) regulate light propagation and tissue heating in corals. Measurement-informed simulations of coupled radiative and heat transfer showed that high host pigment concentrations increase broadband scattering while strongly absorbing light within their characteristic spectral bands, reducing light penetration into deeper tissue layers and producing localized energy deposition. Under high symbiont densities, tissue heating was governed by absorption by both symbionts and host pigments, whereas in bleached corals, host pigments became the dominant source of heat generation. Heat generated by CP absorption was concentrated within the epidermis, while symbiont absorption caused deeper heating within the gastrodermis. Unlike CPs, FPs additionally redistributed a fraction of the absorbed photons through lower energy fluorescence emission, increasing scalar irradiance at longer wavelengths and shifting energy deposition deeper into the tissue, thereby enhancing subsurface heating relative to the corresponding non-fluorescent CP scenario. In the model, this pathway changes scalar irradiance and predicted energy deposition relative to the same FP parameterization with fluorescence disabled. The magnitude of this effect varied with FP concentration, spatial organization (granular or diffuse), and symbiont density. Model outputs were supported by surface light and temperature microsensor measurements across sampled coral phenotypes and symbiont densities. Together, our results demonstrate distinct modeled optical pathways for CPs and FPs that regulate not only light propagation but also the spatial distribution of heat generation within coral tissues. By balancing light attenuation, spectral redistribution, and heat generation, host pigmentation is thus likely to influence photoacclimation, bleaching susceptibility, and recovery of the coral-algal symbiosis.

## 1. Introduction

Coral reefs are among the most productive and biologically diverse ecosystems on Earth, despite their limited distribution in mostly oligotrophic, tropical waters.^2,5^ Their ecological success is fundamentally driven by a mutualistic symbiosis between the coral host and intracellular photosynthetic dinoflagellates in the family Symbiodiniaceae, which provide the majority of the host’s nutrition via translocation of photosynthates.^2,5^ Light is therefore the primary energy source sustaining coral reef productivity, regulating photosynthesis, and stimulating calcification. ^2,5^ At the same time, excess irradiance can cause photodamage and, particularly when combined with elevated temperatures, destabilize the coral-algal symbiosis, thereby exacerbating bleaching and contributing to reef degradation.^8–10^

Corals are efficient natural light-harvesting systems, where both skeletal scattering and tissue optical properties jointly control the internal light field experienced by symbionts.^11^ The interaction of incident light with coral tissue and skeleton creates a highly complex optical microenvironment, governed by multiple scattering, absorption, and wavelength-dependent redistribution processes.^1,6,7,11,12^ The highly scattering aragonite skeleton acts as a diffuse reflector, increasing photon pathlengths and enhancing scalar irradiance within the overlying tissue, thereby improving light absorption efficiency by symbionts.^13^ At the same time, coral tissue itself contributes significantly to light redistribution through scattering and absorption processes, generating steep vertical and lateral gradients in scalar irradiance and creating distinct optical microniches within the tissue. ^7^ These gradients can result in orders-of-magnitude variation in light availability over sub-millimeter scales, exposing symbionts to highly heterogeneous light environments. ^7^

Light absorption within coral tissues is closely coupled to thermal dynamics, as a substantial fraction of absorbed radiant energy is dissipated as heat.^14,15^ A major component of this heat production can arise from non-photochemical quenching (NPQ) by the symbiotic algae, whereby excess absorbed light energy is dissipated as heat to protect the photosynthetic apparatus under high irradiance conditions.^10^ Tissue heating leads to the formation of temperature gradients across the tissue-water interface and within the coral, particularly under high irradiance and low flow conditions.^14–16^ Heat transport is governed by the interplay between radiative energy deposition, conductive transfer within tissue and skeleton, and convective exchange across the thermal boundary layer.^12,17^ Localized heating can exacerbate photophysiological stress, contributing to photoinhibition and bleaching.^10,18^ Importantly, the efficiency of heat dissipation is strongly influenced by microscale hydrodynamics, optical properties and morphology, linking optical and thermal processes in determining coral stress responses.^12^

In addition to their colony morphology, corals actively regulate their internal light environment through the expression of host-derived pigments, primarily green fluorescent protein (GFP)-like pigments.^19–22^ These pigments include both fluorescent proteins (FPs), which absorb light and re-emit it at longer wavelengths, and non-fluorescent chromoproteins (CPs), which primarily act as strong absorbers.^19,20,23^ Host pigments can constitute a significant fraction of coral tissue protein content and exhibit diverse spectral properties spanning much of the visible range.^19,20,24,25^ Their spatial distribution and dynamic regulation enable corals to modulate light availability to symbionts under varying environmental conditions.^22,26–29^

Mechanistically, host pigments influence light transport through multiple pathways. CPs attenuate incident radiation via strong absorption, thereby reducing internal light levels and providing photoprotection under high irradiance.^22,28,29^ In contrast, fluorescent proteins introduce wavelength conversion pathways by absorbing high-energy light and re-emitting it at longer, less energetic wavelengths, redistributing light to deeper tissue layers.^21,23,27^ Additionally, interactions between pigments, including radiative transfer and Förster resonance energy transfer (FRET), may further modify the spectral and spatial distribution of light within the tissue.^19,30^ FRET is non-radiative, short-range transfer of excitation between donor and acceptor pigments; it is distinct from emitted-photon transport^19,30,31^ and is not represented in the model discussed in this work. The net effect of these processes can be either photoprotective or photoenhancing, depending on environmental conditions and pigment expression levels.^21,26,29^

Host pigment regulation is particularly important during stress events such as coral bleaching.^21,32^ During bleaching, the loss of symbionts and photosynthetic pigments leads to a substantial increase in internal light levels due to reduced absorption and enhanced backscattering from the skeleton, creating a positive optical feedback loop that exacerbates photodamage.^33^ In response, corals can increase the expression of FPs and CPs, which modify the internal light environment and can reduce excess irradiance reaching deeper tissue layers, thereby promoting conditions favorable for symbiont recovery and recolonization.^28^

Despite significant advances in experimental characterization of coral optics^4,7,11,26^, a quantitative and predictive understanding of the coupled radiative, thermal, and physiological processes is still lacking. Numerical modeling approaches, including Monte Carlo simulations, radiative transfer models, and finite element computational methods, have been widely used to simulate light propagation in corals and other turbid biological media.^1,12,34,35^ These models capture the effects of multiple scattering, absorption, and anisotropy on photon transport and have demonstrated the importance of skeletal reflectance and tissue optical properties in shaping internal light fields.^1,12,34,35^ More recent multiphysics frameworks integrate light transport with heat transfer and mass exchange processes in realistic three-dimensional coral geometries, enabling the investigation of coupled light, temperature and oxygen dynamics at the colony to sub-tissue scale.^12^ However, most existing models simplify or neglect host pigment dynamics, particularly fluorescence emission and pigment-specific light absorption/scattering, leaving their role in shaping the coupled light and thermal microenvironment insufficiently resolved. Bridging this gap requires integrating experimental determination of coral optical properties with multiphysics simulations that capture their effects on light redistribution, energy dissipation, and heat generation. In this study, we combine optical and thermal microsensor measurements with mechanistic modeling to investigate how CP- and FP-rich coral tissue modulates light transport and thermal dynamics. By explicitly incorporating host pigment and symbiont density-dependent optical properties, including scattering, absorption and fluorescence, we resolve how different host pigment configurations influence depth-resolved light and temperature fields in coral tissue. This approach provides a mechanistic understanding of how coral host pigments regulate the optical and thermal microenvironment experienced by both the coral host and its symbiotic algae.

## 3. Results and discussion

To resolve how coral host pigments modify light transport and heat generation, we combined optical and thermal microsensor measurements with mechanistic modeling incorporating pigment- and symbiont-density-dependent optical properties, including wavelength-specific scattering, absorption, and fluorescence.

Figure 1 summarizes the conceptual framework and model geometry used to interpret the measurements. A representative coral fragment with high symbiont density is shown in Fig. 1A. The layered coral tissue architecture implemented in the model is illustrated in Fig. 1B, comprising the mucus layer, epidermis (containing host pigments), mesoglea, gastrodermis (containing symbionts), gastrovascular cavity, and the underlying calcium carbonate skeleton (Fig. S5A). The skeleton and the calicoblast are combined, since both layers were assumed to be dominated by porous aragonite (CaCO_3_).

**Figure 1.**
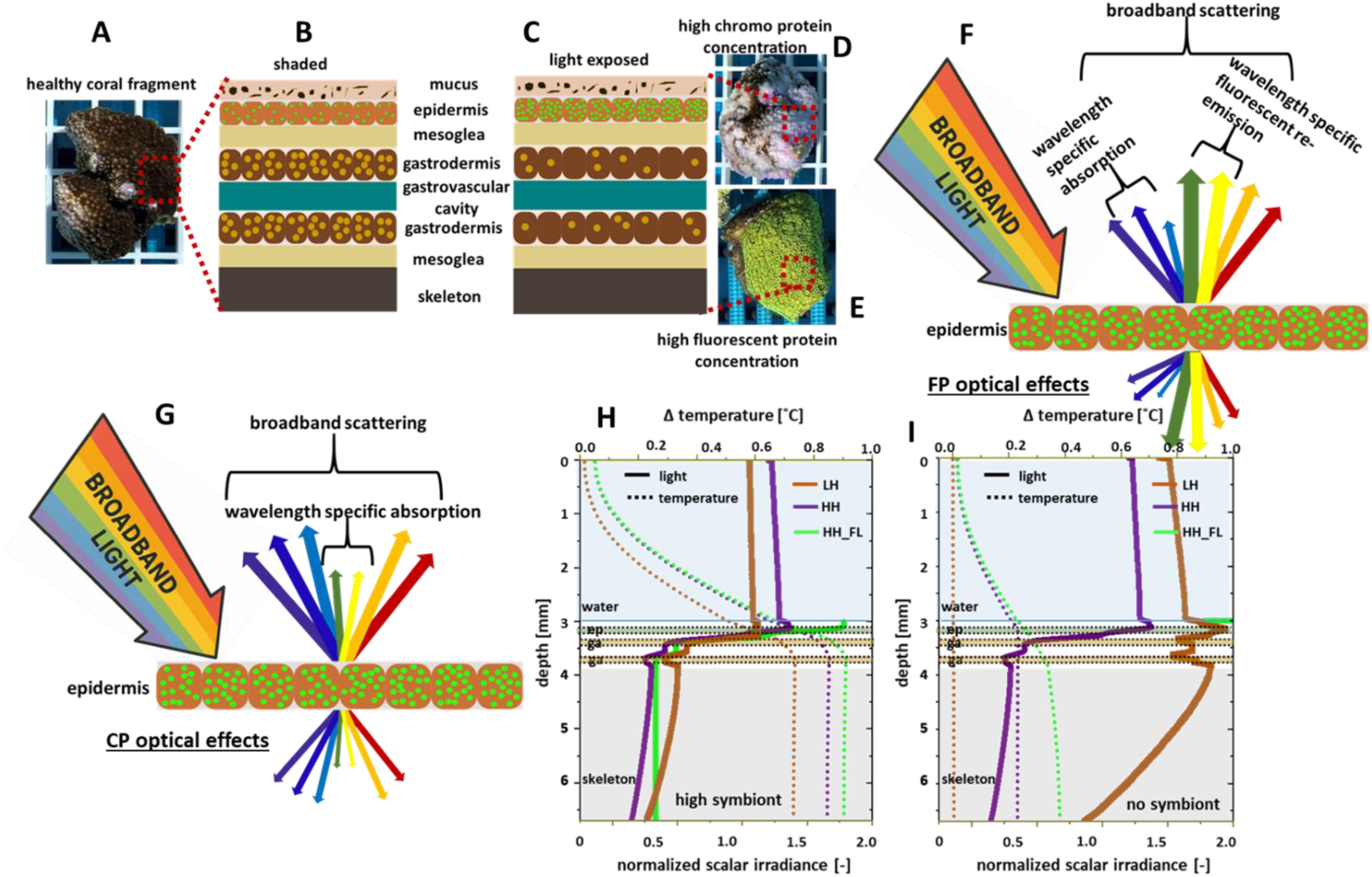
Conceptual framework showing effects of host pigment and symbiont concentration on light transport and heat generation in coral tissue. (A) Photograph of coral fragment, of Montipora sp., with high symbiont density, sampled from a shaded region. (B) Schematic representation of the internal architecture in healthy shaded coral tissue, including mucus, epidermis containing coral host pigments, mesoglea, gastrodermis with symbiotic dinoflagellates, gastrovascular cavity, and the underlying skeleton. (C) Schematic of the tissue structure for a high-light exposed fragment, characterized by reduced symbiont density and increased host pigment concentrations. (D) Bleached coral fragment, of Montipora sp., exhibiting high chromoprotein (CP) concentration. (E) Coral fragment of Cyphastrea microphthalma, sampled from a high-light region, showing high fluorescent protein (FP) concentration. (F) Conceptual illustration of broadband light interaction with fluorescent proteins (FP). (G) Conceptual illustration of optical effects of chromoproteins (CP). (H) Simulated depth profiles of normalized scalar irradiance (PAR) and temperature (dotted lines) for coral with high symbiont concentrations, comparing low-host-pigment concentration (LH) with high-host-pigment concentration with fluorescence disabled (HH) or enabled (HH_FL). (I) Same comparison in the no-symbiont model. LH: fragment with low host protein concentration; HH: fragment with high host protein concentration, with no HP fluorescence; HH_FL: fragment with high host protein concentration, with HP fluorescence.

The framework focuses on how variations in symbiont abundance and host pigment concentration influence light propagation and heat generation within coral tissue. Corals with reduced symbiont density and elevated concentrations of host pigments (Fig. 1C) exhibit substantially different optical environments compared to non-pigmented or weakly pigmented ones. Different pigmentation patterns in corals can arise from multiple biological processes, including photoacclimation (as observed for example on the light-exposed vs shaded regions of pigmented colonies^36^), colorful bleaching^32^, or genetic polymorphism (i.e. colour morphs with different genetic potential to produce host pigments)^37^. The model was designed to evaluate the optical and thermal consequences of host pigments independent of the specific biological processes responsible for their accumulation.

Two characteristic host pigment classes are illustrated in Fig. 1D–E, corresponding to chromoproteins (CPs) and fluorescent proteins (FPs), respectively. These pigments differ fundamentally in their photophysical properties. FPs absorb incident light within defined excitation bands and re-emit part of the absorbed energy as longer-wavelength fluorescence, while exhibiting broadband elastic scattering across the remaining PAR spectrum^20,38^ (Fig. 1F). The emitted photons contribute to the internal light field through subsequent scattering and absorption^26,27^. In contrast, CPs are non-fluorescent pigments that primarily influence the internal light field through strong broadband scattering combined with wavelength-specific absorption (Fig. 1G).^22,29^

The simulated scalar irradiance profiles (example Fig 1H) exhibit several local maxima and minima arising from the interplay between absorption and scattering within the different tissue layers. In host pigment-rich epidermal layers, scalar irradiance can increase locally due to enhanced scattering, which redirects photons and increases their residence time within the tissue. In contrast, minima occur within the gastrodermal layers owing to strong absorption by symbiont pigments. A modest increase in scalar irradiance is observed within the intervening gastric cavity due to its very low absorption, allowing photons transmitted and scattered by the surrounding tissue and skeleton to persist before encountering the lower gastrodermis. A further increase occurs near the tissue–skeleton interface due to strong skeletal backscattering, which redirects photons back into the overlying tissue and enhances photon pathlengths. Consequently, scalar irradiance does not necessarily decrease exponentially with depth but instead reflects the balance between absorption, scattering, and skeletal reflection throughout the tissue-skeleton system.

Figure 1H, I and S1 conceptually summarize how high host pigment concentrations regulate the coral light and thermal microenvironment under contrasting symbiont densities, illustrating how chromoproteins (CPs) and fluorescent proteins (FPs) differentially modulate internal light propagation and tissue heating. Panel H represents corals with high symbiont and host pigment concentrations, where light absorption and scattering by both symbionts and host pigments jointly shape the internal light field and temperature distribution. Panel I represents bleached corals with very low symbiont density but the same high host pigment concentration (HH) as in panel H. Under these conditions, the internal light field and heat generation are primarily governed by host pigments and light scattering by the coral skeleton. In both panels, the simulated profiles are compared with those for corals with low host pigment concentrations (LH) at the corresponding symbiont density. The panels also illustrate the contrasting optical strategies of CPs and fluorescent proteins FPs by comparing simulations in which all absorbed light is converted to heat (HH), representing CPs, with simulations in which 65% of the absorbed light is re-emitted as fluorescence (quantum yield = 0.65)^38^ and the remaining 35% is dissipated as heat (HH_FL), representing FPs.

Fig. 1H, I and Fig. S1, conceptually summarize how high host pigment concentrations increase broadband scattering within the epidermis, reducing light penetration into deeper tissue layers^29^, while strongly absorbing light within their characteristic spectral bands^20,23^. This combination of enhanced scattering and spectrally selective absorption results in localized energy deposition and higher tissue temperatures compared with corals containing low host pigment concentrations at the same symbiont density. When symbiont density is high (Fig. 1H), absorption by both symbionts and host pigments contributes to tissue heating. In contrast, in bleached corals with very low symbiont densities (Fig. 1I and Fig. S1), tissue heating associated with symbiont absorption is greatly reduced, and heat generation is dominated by absorption by the host pigments. The thermal effect of host pigments therefore reflects two competing optical processes, where broadband scattering reduces the internal irradiance available for absorption, while strong absorption within narrow spectral bands increases local energy deposition. The balance between these processes determines the net tissue heating, which increases with host pigment concentration.

Fluorescent protein (FP)-rich corals further modify the internal light field through fluorescence-mediated spectral redistribution. Simulations comparing low FP concentration (LH), high FP concentration without fluorescence (HH; equivalent to chromoproteins scenario, CPs), and high FP concentration with fluorescence (HH_FL; equivalent to FPs) show that, at high FP concentrations, fluorescence enhances scalar irradiance and tissue heating relative to the non-fluorescent (CP) scenario (Fig. 1H). This enhancement arises because fluorescence re-emits a fraction of the absorbed photons at longer wavelengths, allowing them to undergo further scattering and absorption within the coral tissue and skeleton. At the same time, a fraction of both the incident and fluorescent photons is lost through elastic scattering out of the tissue, such that the net effect depends on the balance between photon escape and fluorescence-mediated redistribution. The effect was most pronounced in a simulated bleached coral (Fig. 1I) with very high FP concentrations, where the modelled fluorescence emission increased by up to 30-fold relative to the deposited light energy within the FP absorption band owing to the absence of competing symbiont absorption and the high molar extinction coefficient of FPs. Under otherwise identical conditions, this resulted in greater subsurface energy deposition and higher tissue temperatures than in the corresponding CP scenario. This represents a hypothetical high-FP bleached scenario, that likely represents an extreme; its biological plausibility depends on whether these pigment abundances and locations occur in vivo.

Together, these results support distinct optical strategies in CP and FP bearing corals represent distinct optical strategies for regulating light availability and tissue heating within coral tissues. Both pigment classes reduce internal light penetration through enhanced broadband scattering and wavelength-specific absorption. However, unlike CPs, FPs additionally re-emit a fraction of the absorbed energy as fluorescence, introducing a radiative pathway that redistributes light within the tissue. This fluorescence-mediated redistribution partially offsets the reduction in scalar irradiance caused by elastic scattering, resulting in greater subsurface light availability, energy deposition, and tissue heating than in the corresponding CP scenario. In both cases, absorbed photon energy is dissipated as heat through non-radiative relaxation processes^39^, whereas FPs partition the absorbed energy between heat generation and fluorescence emission. These contrasting optical strategies are likely to influence how corals regulate their internal optical and thermal microenvironment under both stressful and non-stressful environmental conditions. By reducing the amount of excess light penetrating deeper tissue layers, high host pigment concentrations may also contribute to photoprotection of the remaining symbionts in stressed corals.^21,40^ This modulation of the internal light environment could create more favorable conditions for symbiont survival and recolonization during recovery from bleaching. Such a mechanism is consistent with previous observations that bleached corals expressing high concentrations of chromoproteins recover more rapidly than non-colourful morphs.^28^ However, our simulations show that this photoprotective effect may come at the cost of a modest increase in internal tissue temperature, which could remain tolerable under mild heat stress.

To further examine these optical strategies, we present four representative examples combining measurements and simulations from coral fragments collected on the Great Barrier Reef (except *A. curvata* example 4, aquarium-reared). Two examples focus on CP-rich corals and the effects of CPs on light scattering and heat deposition, whereas two additional examples investigate FP-rich corals and fluorescence-mediated modification of the internal light and thermal microenvironment.

### 3.1 Chromoprotein-rich phenotypes

To investigate how CP-rich phenotypes regulate light availability and tissue heating, we analyzed two coral species collected from the shallow reef-flat (<2 m depth) at Heron Island, Great Barrier Reef, Australia. In *Isopora palifera*, a single fragment exhibited strong spatial heterogeneity in CP and symbiont density, with a heavily bleached, CP-containing and light-exposed surface area, and a shaded tissue area with high symbiont content (Fig. 2). *Montipora capricornis* fragments were collected from shaded and light-exposed regions of the same colony to assess how local light history regulates CP expression and the resulting optical and thermal microenvironment (Fig. 3). In both cases, fluorescent protein (FP) expression was negligible, as confirmed by weak fluorescence under blue/violet excitation and reflectance measurements. Thus, the observed optical and thermal responses can be attributed primarily to variations in CP concentration and symbiont density.

**Figure 2.**
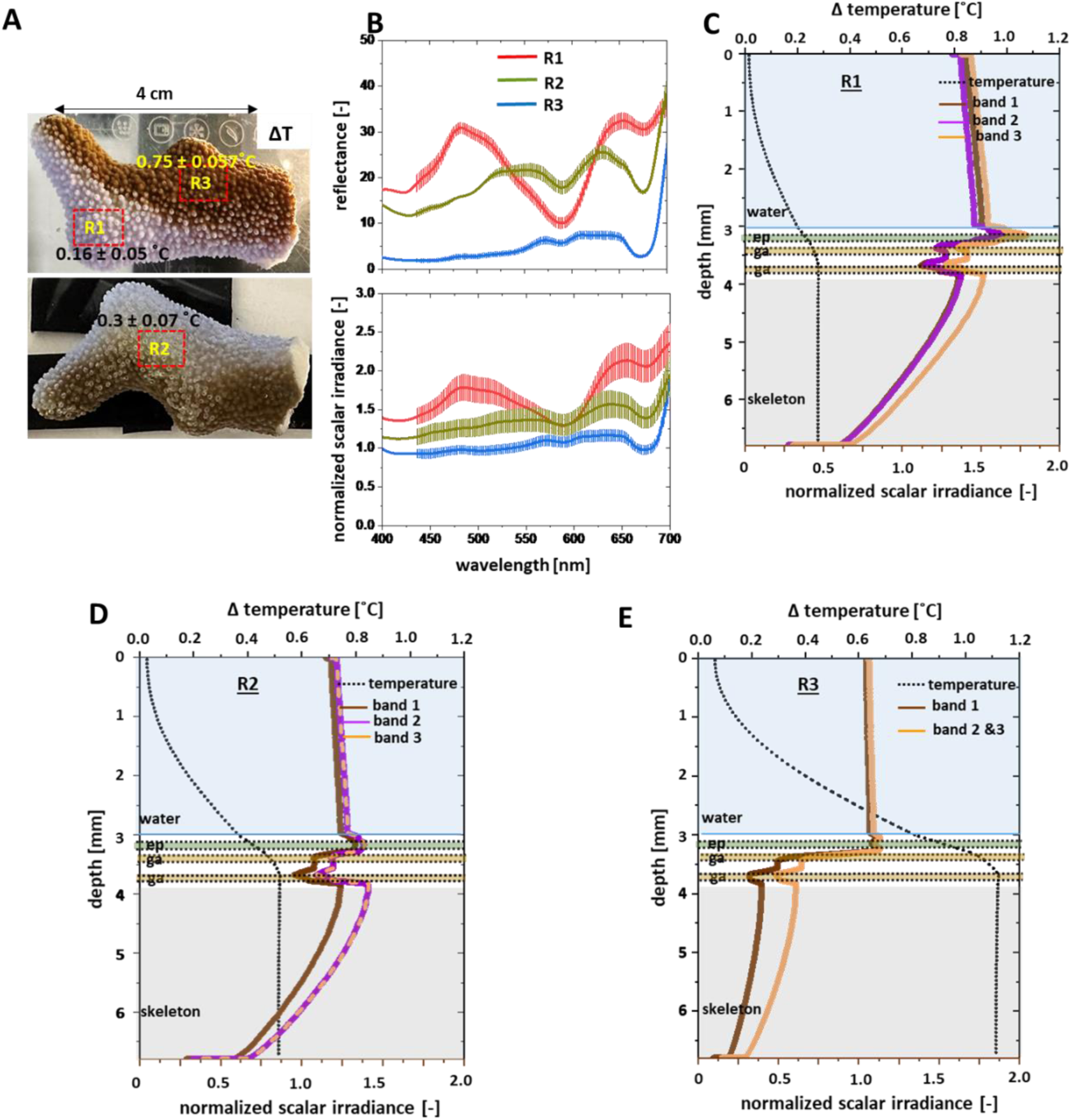
Optical/thermal characterization and model-based analysis of colorful bleaching in an Isopora palifera coral fragment. (A) Photographs of the fragment illustrating spatial heterogeneity in pigmentation and bleaching state. Dashed boxes indicate the regions of interest used for measurements and simulations. R1 corresponds to a strongly bleached region with high chromoprotein concentration, R2 represents a less bleached region with moderate chromoprotein expression, and R3 represents a region with high symbiont density and low chromoprotein concentration. The measured surface temperature increase relative to the overlying water (ΔT) under a downwelling irradiance (1600 µmol photons m⁻² s⁻¹ / 350 W m⁻²) is shown for each region. Lower panel shows the 3D texture scan of the fragment. (B) Optical measurements: reflectance spectra measured from three regions of interest (R1–R3) on the coral fragment. Scalar irradiance spectra measured at the coral tissue-water interface and normalized to the incident irradiance. Error bars represent standard deviation from replicate measurements, n = 6 for each region. (C– E) Simulated depth profiles of normalized scalar irradiance and temperature for regions R1–R3, respectively. Band 1: 400–490 nm and 650–680 nm, principal absorption maxima of Symbiodiniaceae pigments; Band 2: 540–620 nm, absorption range for CP; band 3: remaining wavelengths in the PAR range (400-700 nm); ep: epidermis; ga: gastrodermis.

**Figure 3.**
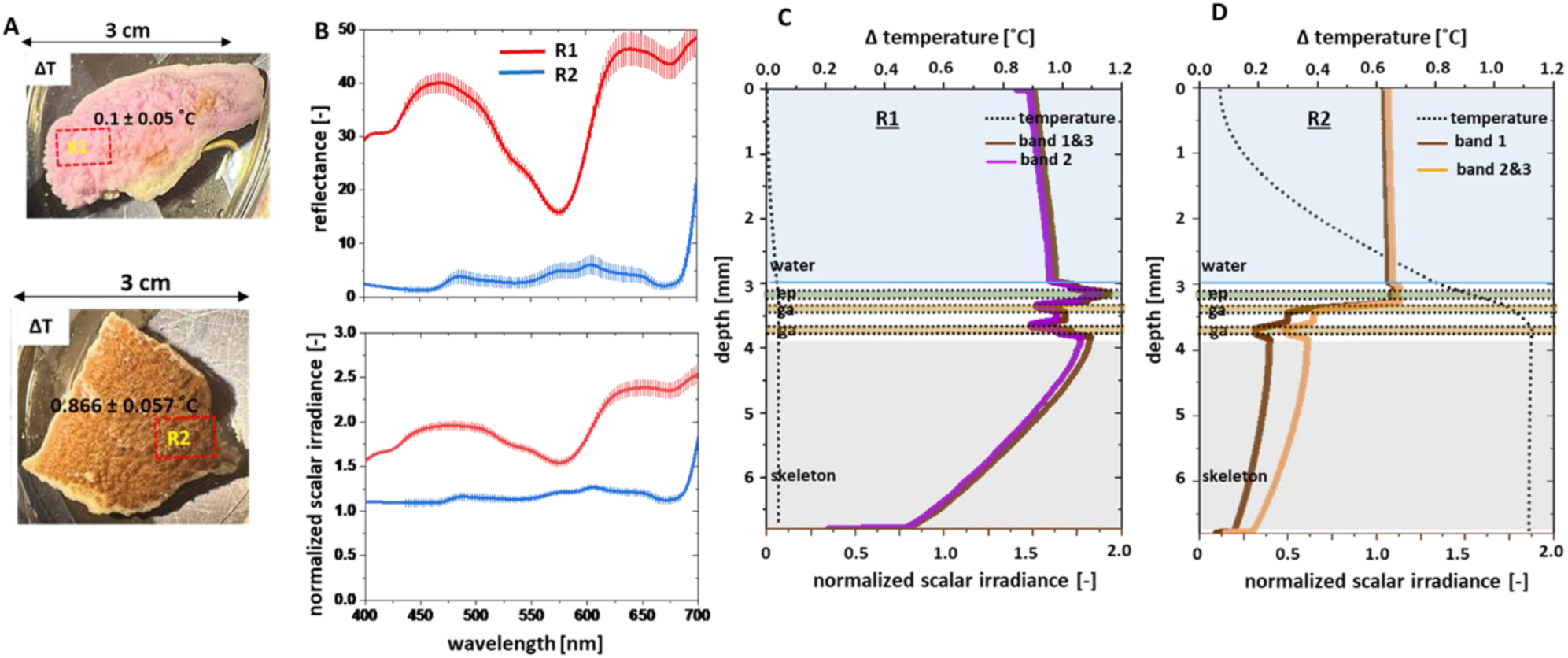
Optical/thermal characterization and model-based analysis of colorful bleaching in Montipora capricornis fragments collected from contrasting regions of the same colony. (A) Photographs of coral fragments showing distinct pigmentation states. R1: fragment from a high-light-exposed region, exhibiting bleaching with high CP concentration and very low symbiont density (pink appearance); R2: fragment from a shaded region with lower CP expression and much higher symbiont density (brown appearance). Insets indicate regions of interest used for temperature measurements. Corresponding surface temperature increases (ΔT) under downwelling irradiance (1600 µmol photons m⁻² s⁻¹ / 350 W m⁻²) are shown for each region. (B) Reflectance spectra (top) and normalized scalar irradiance (bottom) for R1 and R2. (C–D) Simulated depth profiles of normalized scalar irradiance (bands 1–3) and temperature for R1 and R2, respectively. Band 1: 400–490 nm and 650–680 nm, principal absorption maxima of Symbiodiniaceae pigments; Band 2: 540–620 nm, absorption range, for CP; band 3: rest; ep: epidermis; ga: gastrodermis. For R1, wavebands 1 and 3 are identical because a fully bleached state was assumed. For R2, wavebands 2 and 3 are identical because CP absorption is absent.

#### 3.1.1 Isopora palifera

The coral fragment exhibited pronounced spatial heterogeneity in host pigment concentration and symbiont density (Fig. 2A). Three regions were identified: strongly bleached tissue with high CP concentration (R1), partially bleached tissue with moderate CP levels (R2), and a region with higher symbiont density and low CP expression (R3). These differences were reflected in the reflectance spectra (Fig. 2B), where R1 and R2 showed strong absorption in the CP spectral band (∼540–620 nm) and reduced chlorophyll absorption, consistent with lower symbiont densities. In contrast, R3 exhibited weaker CP absorption and more pronounced chlorophyll absorption features, indicating higher symbiont abundance.

Scalar irradiance measurements at the tissue-water interface further revealed how CP expression modifies the surface light field (Fig. 2B). Regions with elevated CP (R1, R2) showed reduced normalized scalar irradiance within the CP absorption band. In the chlorophyll absorption bands, R2 exhibited a stronger attenuation than R1, consistent with its relatively higher symbiont density. Outside these absorption bands, scalar irradiance was enhanced (up to ∼1.25–2× relative to the incident light) due to multiple scattering within the tissue and skeleton. However, overall scalar irradiance across most of the PAR spectrum (excluding the CP absorption band) remained higher in R1 than in R2, reflecting its lower symbiont density and thus reduced light absorption. In contrast, R3 exhibited lower normalized scalar irradiance across the PAR range (≈1), consistent with stronger light absorption by symbionts.

These differences in light availability were directly reflected in tissue heating under high irradiance (1600 µmol photons m⁻² s⁻¹; Fig. 2A), as measured surface temperature increased. The symbiont-rich region R3 showed the highest temperature increase (ΔT = 0.75 ± 0.05 °C), while R2 and R1 exhibited intermediate (ΔT = 0.30 ± 0.07 °C) and minimal heating (ΔT = 0.16 ± 0.05 °C), respectively. This trend indicates that lower CP expression and higher symbiont density promote deeper light penetration and absorption, resulting in increased heat generation.

Simulations (Fig. 2C–E) provide a mechanistic explanation for these observations. The optical properties and model parameters used in the simulations, inferred from the reflectance profiles, are summarized in Tables 1–3. The simulated light fields and temperature profiles are consistent with the measured surface scalar irradiance and temperature patterns:

**Table 1.**
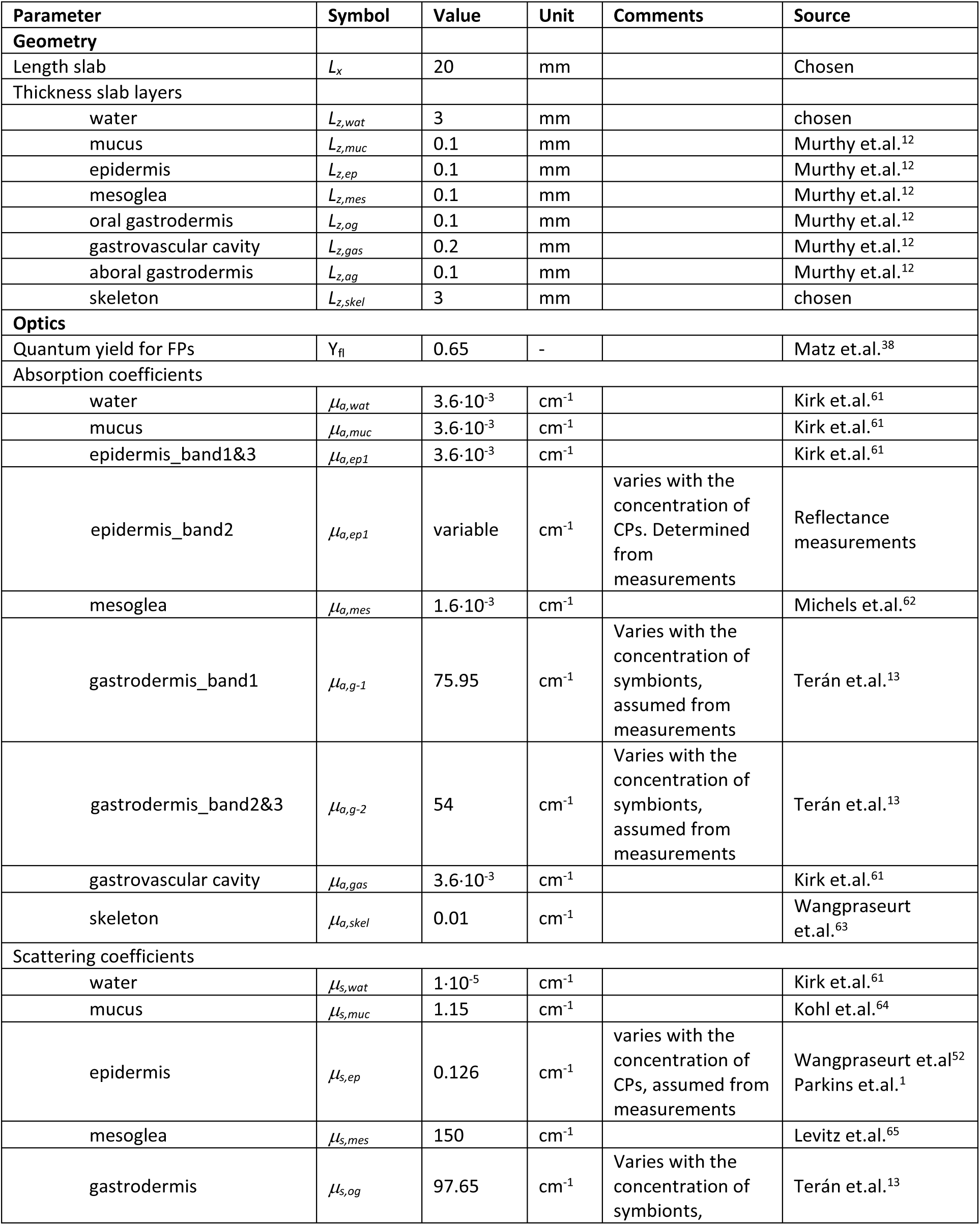
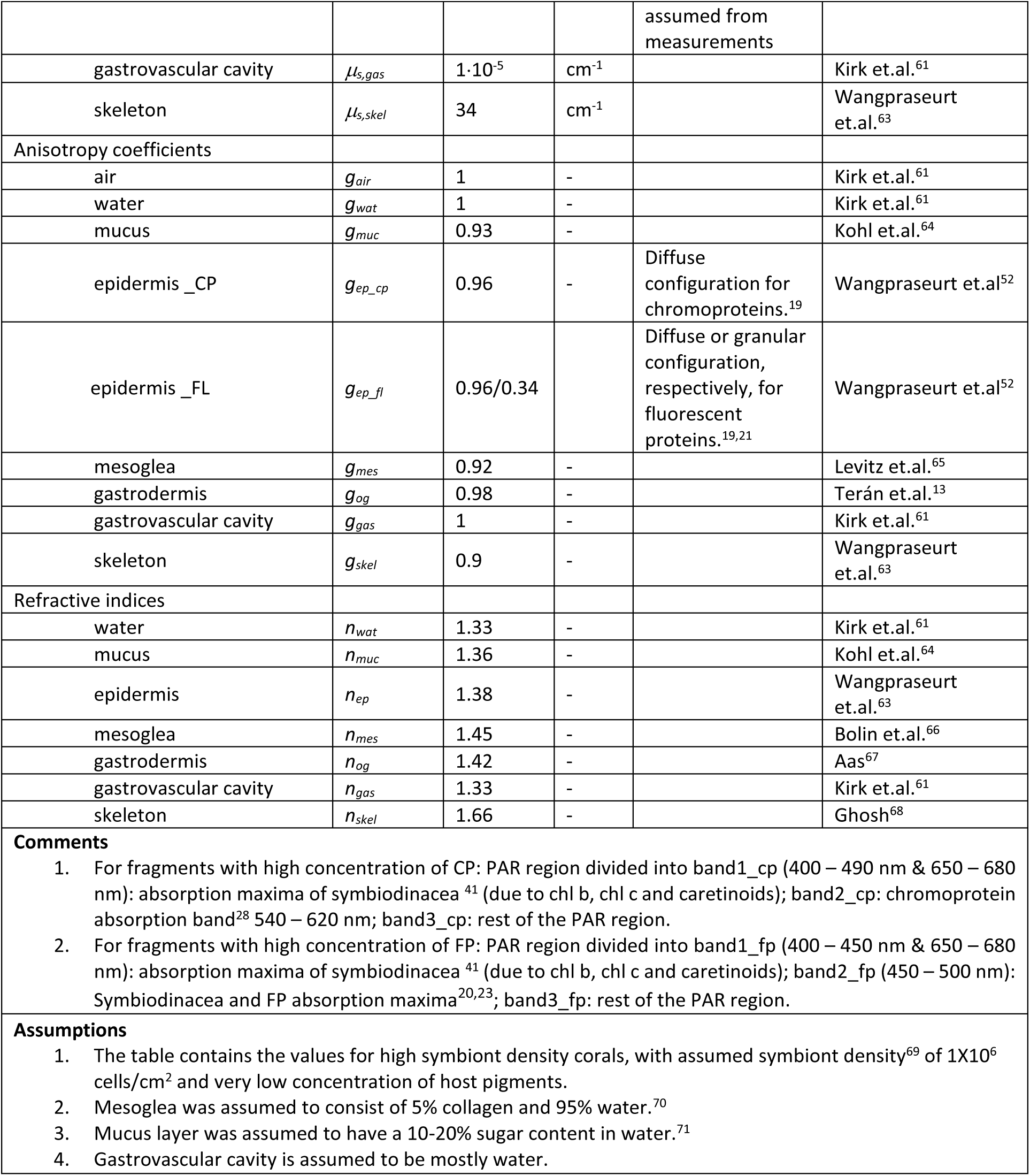
Model parameters for coral geometry and optics.

**Table 2.** Model parameters for flow, heat, mass transport and biokinetics.

| Parameter | Symbol | Value | Unit | Source |
| --- | --- | --- | --- | --- |
| **Flow and heat** |  |  |  |  |
| Viscosity water | $\eta_{wat}$ | 0.0009 | Pa s | Incropera et.al.<sup>72</sup> |
| **Densities** |  |  |  |  |
| water | $\rho_{wat}$ | 997 | kg m<sup>-3</sup> | Incropera et.al.<sup>72</sup> |
| tissue (all) | $\rho_{tis}$ | 1109 | kg m<sup>-3</sup> | Hasgall et.al.<sup>73</sup> |
| skeleton | $\rho_{skel}$ | 2930 | kg m<sup>-3</sup> | Bragg <sup>74</sup> |
| Heat conductivities |  |  |  |  |
| water | $k_{wat}$ | 0.6 | W m<sup>-1</sup> K<sup>-1</sup> | Incropera et.al.<sup>72</sup> |
| tissue (all) | $k_{tis}$ | 0.37 | W m<sup>-1</sup> K<sup>-1</sup> | Hasgall et.al.<sup>73</sup> |
| skeleton | $k_{skel}$ | 1.5 | W m<sup>-1</sup> K<sup>-1</sup> | Jimenez et.al.<sup>16</sup> |
| Heat capacities |  |  |  |  |
| water | $C_{p,wat}$ | 4183 | J kg<sup>-1</sup> K<sup>-1</sup> | Incropera et.al.<sup>72</sup> |
| tissue (all) | $C_{p,tis}$ | 3391 | J kg<sup>-1</sup> K<sup>-1</sup> | Hasgall et.al.<sup>73</sup> |
| skeleton | $C_{p,skel}$ | 811 | J kg<sup>-1</sup> K<sup>-1</sup> | Staveley et.al.<sup>75</sup> |
| **Experimental parameters** |  |  |  |  |
| Average water velocity | $v_0$ | 1 | mm s<sup>-1</sup> | measurement |
| Ambient temperature | $T_0$ | varies | °C | measurement |
| Incident downwelling photon irradiance | $I_{id}$ | varies | μmol photon m<sup>-2</sup> s<sup>-1</sup> | measurement |

**Table 3:**
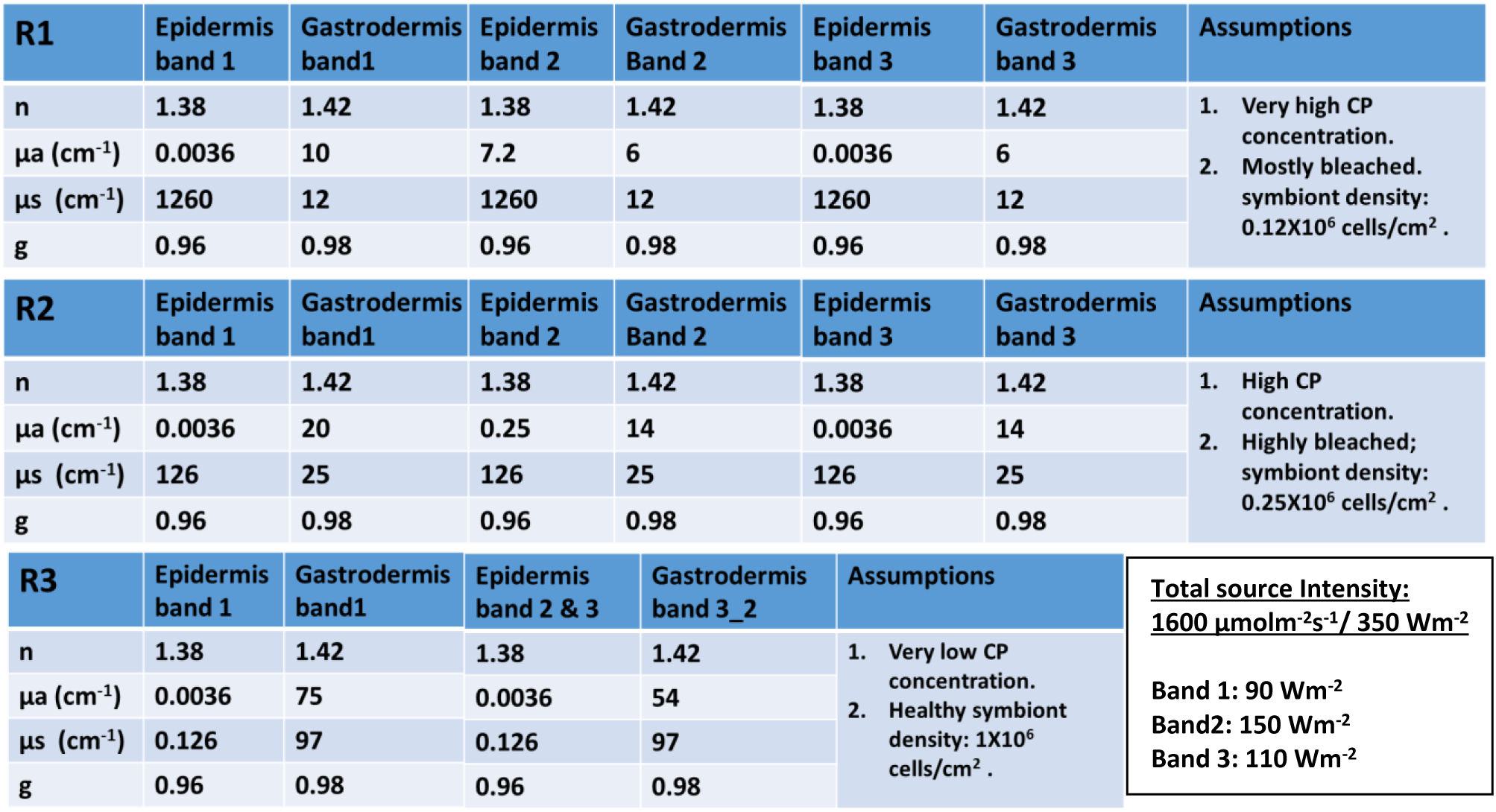
Summary of the epidermal and gastrodermal optical properties of the Isopora palifera fragment, divided into three distinct zones with different symbiont and CP concentrations as indicated by reflectance spectra and visual inspection. R1: mostly bleached region with very high CP concentration; R2: highly bleached region with high CP concentration; R3: shaded (from light) region with healthy symbiont density and very low CP concentration. Optical properties were determined as described in our previous work^1,2^. For the colorfully bleached regions, the epidermal absorption coefficient (µ_a_) was calculated from the reflectance spectra.^4,5^ µ_s_: scattering coefficient; g: anisotropy of scattering and n: refractive index. The table also indicates the incident light intensity for each waveband in the PAR region. Band 1: 400 – 490 nm & 650 – 680 nm; Band 2: 540 – 620 nm; Band 3: rest of the PAR region.

| R1 | Epidermis band 1 | Gastrodermis band1 | Epidermis band 2 | Gastrodermis Band 2 | Epidermis band 3 | Gastrodermis band 3 | Assumptions |
| --- | --- | --- | --- | --- | --- | --- | --- |
| n | 1.38 | 1.42 | 1.38 | 1.42 | 1.38 | 1.42 | 1. Very high CP concentration.<br>2. Mostly bleached. symbiont density: 0.12X10<sup>6</sup> cells/cm<sup>2</sup> . |
| μa (cm<sup>-1</sup>) | 0.0036 | 10 | 7.2 | 6 | 0.0036 | 6 |  |
| μs (cm<sup>-1</sup>) | 1260 | 12 | 1260 | 12 | 1260 | 12 |  |
| g | 0.96 | 0.98 | 0.96 | 0.98 | 0.96 | 0.98 |  |
| R2 | Epidermis band 1 | Gastrodermis band1 | Epidermis band 2 | Gastrodermis Band 2 | Epidermis band 3 | Gastrodermis band 3 | Assumptions |
| n | 1.38 | 1.42 | 1.38 | 1.42 | 1.38 | 1.42 | 1. High CP concentration.<br>2. Highly bleached; symbiont density: 0.25X10<sup>6</sup> cells/cm<sup>2</sup> . |
| μa (cm<sup>-1</sup>) | 0.0036 | 20 | 0.25 | 14 | 0.0036 | 14 |  |
| μs (cm<sup>-1</sup>) | 126 | 25 | 126 | 25 | 126 | 25 |  |
| g | 0.96 | 0.98 | 0.96 | 0.98 | 0.96 | 0.98 |  |
| R3 | Epidermis band 1 | Gastrodermis band1 | Epidermis band 2 & 3 | Gastrodermis band 3_2 | Assumptions |  | Total source Intensity:<br>1600 μmolm<sup>-2</sup>s<sup>-1</sup>/ 350 Wm<sup>-2</sup><br>Band 1: 90 Wm<sup>-2</sup><br>Band2: 150 Wm<sup>-2</sup><br>Band 3: 110 Wm<sup>-2</sup> |
| n | 1.38 | 1.42 | 1.38 | 1.42 | 1. Very low CP concentration.<br>2. Healthy symbiont density: 1X10<sup>6</sup> cells/cm<sup>2</sup> . |  |  |
| μa (cm<sup>-1</sup>) | 0.0036 | 75 | 0.0036 | 54 |  |  |  |
| μs (cm<sup>-1</sup>) | 0.126 | 97 | 0.126 | 97 |  |  |  |
| g | 0.96 | 0.98 | 0.96 | 0.98 |  |  |  |

Across regions R1–R3, the thermal response reflected the combined effects of CP concentration, symbiont density, and light penetration. In R1 and R2, elevated CP concentrations enhanced broadband scattering within the epidermis across wavebands 1–3, thereby reducing the penetration of incident light into deeper tissue layers. CP absorption within waveband 2 was largely confined to the epidermis and contributed to localized energy deposition and heating. However, the accompanying reduction in scalar irradiance limits energy deposition at depth. R1 had the highest CP concentration and the lowest symbiont density. Although CP absorption produces localized epidermal heating, the very low symbiont density substantially limited absorption and heat generation in deeper tissue layers, resulting in the smallest temperature increase. R2 had a lower CP concentration but a higher symbiont density than R1. Stronger gastrodermal absorption by symbionts therefore produced an intermediate temperature increase. In contrast, R3 had the lowest CP concentration and the highest symbiont density. Reduced CP-mediated scattering allows more light to penetrate to the gastrodermis, where it is efficiently absorbed by the abundant symbionts. Because approximately 99% of the absorbed energy^15^ is converted to heat in the gastrodermis, R3 exhibited the largest temperature increase, followed by R2 and then R1, consistent with the experimental measurements. Thus, the overall thermal response results from the combined effects of CP and symbiont absorption, with the magnitude and spatial distribution of tissue heating determined by the concentrations of both CPs and symbionts. Heat generated by CPs occurs near the tissue surface and is more readily dissipated to the surrounding seawater, whereas heat generated by symbionts occurs deeper within the tissue, where heat exchange is less efficient.

#### 3.1.2 Montipora capricornis

To further examine the coupled effects of chromoprotein (CP) expression and symbiont density during colorful bleaching, we analyzed fragments originating from different regions within one colony of *M. capricornis*: a fragment from the high-light-exposed top of the colony (R1) exhibited strong bleaching and high CP concentration, whereas a fragment from the more shaded part of the colony (R2) retained high symbiont density and low CP expression.

These differences were evident visually and spectrally (Fig. 3A–B, respectively). R1 showed a characteristic pinkish coloration associated with CP-rich bleached tissue, while R2 appeared brown due to abundant symbionts. Reflectance spectra confirmed strong CP absorption in R1 (∼540–620 nm) along with weak chlorophyll features, consistent with near-complete symbiont loss. In contrast, R2 showed no detectable CP absorption and low reflectance across the PAR region, indicating strong light absorption by symbionts.

Scalar irradiance measurements at the tissue–water interface further highlighted these differences (Fig. 3B). In R1, scalar irradiance was strongly enhanced across the visible spectrum (up to ∼2–2.5× relative to the incident light) due to multiple scattering and low symbiont absorption, however this effect was much lower within the CP absorption band (∼1.5x). In contrast, R2 exhibited lower normalized scalar irradiance across the PAR range (≈1), reflecting efficient light absorption by the dense symbiont population.

These optical differences translated into contrasting thermal responses under high irradiance (1600 µmol photons m⁻² s⁻¹ / 350 W m⁻²; Fig. 3A). The CP-rich, symbiont-poor region (R1) showed only a small surface temperature increase (ΔT ≈ 0.1 ± 0.05 °C), whereas the symbiont-rich shaded region (R2) exhibited substantially higher heating (ΔT ≈ 0.87 ± 0.06 °C), highlighting the dominant role of symbiont absorption in heat generation within coral tissue.

The simulations (Fig. 3C–D) provided mechanistic insight into these observations. The optical properties and model parameters used in the simulations are summarized in Tables 1, 2 and 4. The simulated light fields and temperature profiles were consistent with and supported the experimental surface scalar irradiance and temperature measurements. In R1, elevated CP concentrations enhanced broadband scattering within the epidermis (wavebands 1–3; representing symbiont absorption, CP absorption, and remaining PAR wavelengths, respectively), which limits the penetration of incident light into deeper tissue layers. CP absorption within waveband 2 was confined to the epidermis and resulted in more localized energy deposition and heating. However, the accompanying reduction in scalar irradiance, together with the very low (practically zero) symbiont density, substantially limited energy deposition in the gastrodermis, resulting in only a small overall temperature increase.

**Table 4:** Summary of the epidermal and gastrodermal optical properties of the two **Montipora capricornis** fragments, with different symbiont and CP concentrations as indicated by reflectance spectra and visual inspection. R1: Completely bleached fragment with very high CP concentration; R2: shaded (from light) fragment with healthy symbiont density and very low CP concentration. Optical properties were determined as described in our previous work^1,3^. For the colorfully bleached fragment, the epidermal absorption coefficient (µ_a_) was calculated from the reflectance spectra.^4,5^ µ_s_: scattering coefficient; g: anisotropy of scattering and n: refractive index. The table also indicates the incident light intensity for each waveband in the PAR region. Band 1: 400 – 490 nm & 650 – 680 nm; Band 2: 540 – 620 nm; Band 3: rest of the PAR region.

| R1 | Epidermis<br>Band 1&3 | Gastrodermis<br>Band 1&3 | Epidermis<br>Band 2 | Gastrodermis<br>Band 2 | Assumptions | Total source Intensity:<br>1600 $\mu\text{molm}^{-2}\text{s}^{-1}$/ 350 $\text{Wm}^{-2}$<br>Band 1: 90 $\text{Wm}^{-2}$<br>Band2: 150 $\text{Wm}^{-2}$<br>Band 3: 110 $\text{Wm}^{-2}$ |
| --- | --- | --- | --- | --- | --- | --- |
| n | 1.38 | 1.42 | 1.38 | 1.42 | 1. Very high CP concentration.<br>2. Completely bleached. |  |
| $\mu\text{a}$ ($\text{cm}^{-1}$) | 0.0036 | 0.0036 | 2.1 | 0.0036 | | |
| $\mu\text{s}$ ($\text{cm}^{-1}$) | 1260 | 1.26 | 1260 | 1.26 | | |
| g | 0.96 | 0.98 | 0.96 | 0.98 |  |  |
| R2 | Epidermis<br>band 1 | Gastrodermis<br>band1 | Epidermis<br>band 2 & 3 | Gastrodermis<br>band 2&3 | Assumptions |  |
| n | 1.38 | 1.42 | 1.38 | 1.42 | 1. Very low CP concentration.<br>2. Healthy symbiont density:<br>1X10<sup>6</sup> cells/cm<sup>2</sup> . |  |
| $\mu\text{a}$ ($\text{cm}^{-1}$) | 0.0036 | 75 | 0.0036 | 54 | | |
| $\mu\text{s}$ ($\text{cm}^{-1}$) | 0.126 | 97 | 0.126 | 97 | | |
| g | 0.96 | 0.98 | 0.96 | 0.98 |  |  |

In contrast, R2, characterized by very low HP concentration, allowed deeper propagation of light into the gastrodermis. The high density of symbionts resulted in strong attenuation of scalar irradiance in PAR region, particularly in wave band 1, due to efficient absorption of incident light. Approximately 99% of this absorbed energy was converted to heat in the gastrodermis^15^, leading to increased internal temperatures.

Together, the *I. palifera* and *M. capricornis* examples demonstrate that CP-rich tissues can regulate the internal light environment of corals primarily through enhanced broadband scattering and wavelength-specific absorption. The overall thermal response results from the combined effects of CP and symbiont absorption, where the magnitude and spatial distribution of tissue heating is determined by the concentrations of both host pigments and symbionts. Heat generated by CPs is concentrated within the epidermis, close to the tissue-water interface, where it can be dissipated efficiently into the surrounding seawater, whereas heat generated by symbionts originates deeper within the gastrodermis, where thermal dissipation is less efficient and heat accumulation is therefore greater. Consequently, corals with high CP concentrations and low symbiont densities exhibit lower overall tissue warming than corals with low CP concentrations and abundant symbionts, despite localized epidermal heating by CP absorption. These results illustrate a characteristic benefit of CP synthesis in corals: Reducing internal light availability, while minimizing additional heat generation. Though CP absorption generates localized heat within the epidermis, the overall thermal response is concentration dependent and is determined by the combined contributions of CPs and symbionts.

### 3.2 Fluorescent protein-rich phenotypes

In contrast to CPs, FPs additionally modify the internal light field through wavelength-specific absorption followed by fluorescence emission at longer wavelengths.^21,26^ This process redistributes photons within the tissue, altering both the spectral quality and spatial distribution of light.^21,27^ To examine how FP concentration influences light transport and heat generation, we analyzed *Cyphastrea microphthalma* fragments from different colonies exhibiting highly green fluorescent (CG, *C. microphtalma* green morph) and weak cyan fluorescent (CB, *C. microphtalma* brown morph) phenotypes, collected from both light-exposed and shaded regions within each colony to obtain a range of fluorescence intensities and symbiont concentrations. In addition, an aquarium-reared *Astraeosmilia curvata* fragment with distinct fluorescent and non-fluorescent regions was analyzed. These corals showed strong fluorescence under blue excitation (450–500 nm), with emission in the green range (between 510 –570 nm). The brown CB morph also exhibited weak cyan fluorescence (480–500 nm) under excitation at 440–470 nm. However, this fluorescence was not quantified in the spectra presented here, and its intensity was relatively low compared with the green fluorescence.

#### 3.2.1 Cyphastrea microphthalma

To examine the combined effects of symbiont abundance and FP expression, we compared green-FP-rich (CG) and brown (CB) color morphs, (with weak cyan fluorescence also detected in CB) from shaded and high light-exposed conditions (Fig. 4A). Reflectance spectra of both color phenotypes showed lower reflectance across the PAR region for shaded fragments than light-exposed fragments, indicating higher symbiont density in the shaded fragments (Fig. 4B; Table 5). Within each light regime, the CG morph consistently exhibited higher reflectance than the CB morph, suggesting slightly lower symbiont abundance in the fluorescent morph. In contrast to the CB morph, the CG morph displayed characteristic spectral features of FPs, including a dip in the excitation range (450–500 nm) and enhanced reflectance in the emission range (510–570 nm) ^20^. These FP-associated features were more pronounced in the highlight-exposed CG fragments, consistent with reduced masking by symbiont absorption and potentially higher FP concentration, although the latter could not be verified with the current measurement approach. Based on these inferences regarding FP expression and symbiont abundance, and in the absence of direct measurements of FP concentration, the FP concentration of the light-exposed CG color morph was treated as a fitting parameter in the simulations. Starting from the concentration assumed for the shaded CG morph, it was progressively increased by up to tenfold to reproduce the observed temperature differences.

**Figure 4.**
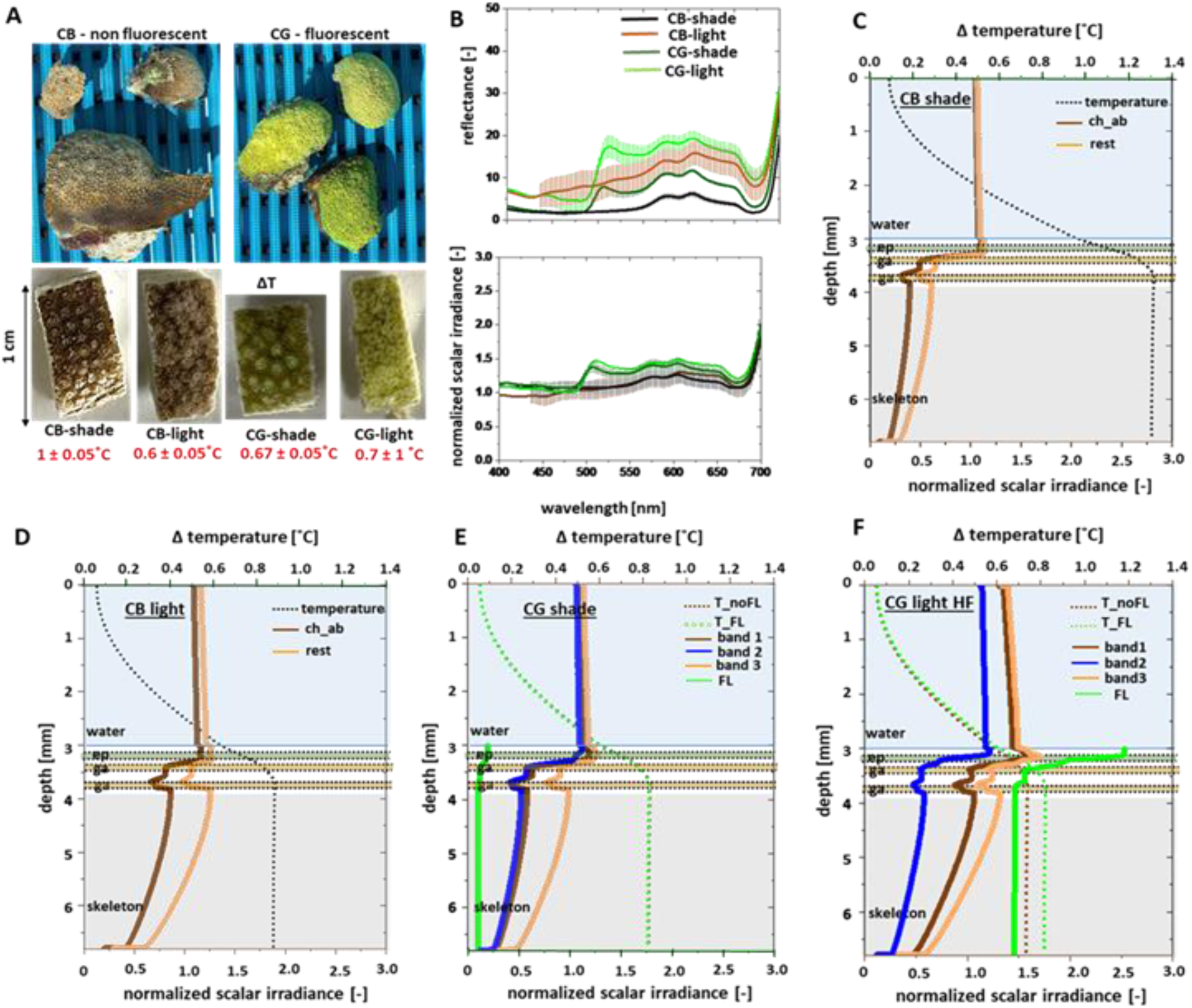
Optical and thermal effects of diffuse fluorescent protein (FP) expression and symbiont density in Cyphastrea microphthalma under contrasting light conditions. (A) Photographs of coral fragments showing brown CB phenotype (weakly fluorescent) and green-FP-rich CG phenotype under white light. Fragments were collected from light-exposed (lower symbiont density) and shaded higher symbiont density) regions. Insets show surface temperature increases (ΔT), w.r.t surrounding water column, under downwelling irradiance (1800 µmol photons m⁻² s⁻¹ / 400 W m⁻²). (B) Reflectance (top) and normalized scalar irradiance (bottom) spectra for CB and CG phenotypes. (C–F) Simulated depth profiles of normalized scalar irradiance (waveband-resolved) and temperature for fragments of CB (non-fluorescent) and CG (fluorescent) morphs collected from shaded and light-exposed parts of the respective colonies. Simulations for CG include fluorescence emission (FL) as a secondary light source, and temperature profiles are shown with (T_FL) and without (T_noFL) fluorescence re-emission, under relatively high FP concentration (HF). Band 1: 400–450 nm and 650–680 nm, principal absorption maxima of Symbiodiniaceae pigments; Band 2: 450–500 nm, absorption range for FP and Symbiodiniaceae pigments; band 3: remaining wavelengths in the PAR range (400-700 nm); ep: epidermis; ga: gastrodermis.

**Table 5:** Summary of the epidermal and gastrodermal optical properties of the four Cyphastrea microphthalma fragments, with different symbiont and FP concentrations as indicated by reflectance spectra and visual inspection. CB-shade: Shaded (from light), non-fluorescent phenotype, fragment with healthy symbiont density and very low FP concentration. CB-light: Light exposed, non-fluorescent phenotype, fragment with partially bleached symbiont density and very low FP concentration. CG-shade: Shaded (from light), fluorescent phenotype, fragment with healthy to partially bleached symbiont density and high FP concentration. CG-light: Shaded (from light), fluorescent phenotype, fragment with mostly symbiont density and very high FP concentration. Optical properties were determined as described in our previous work^1,2,6,7^. The epidermal absorption coefficient (µ_a_) was calculated as described in methods section. µ_s_: scattering coefficient; g: anisotropy of scattering and n: refractive index. The table also indicates the incident light intensity for each waveband in the PAR region. Band 1: 400 – 450 nm & 650 – 680 nm; Band 2: 450 – 500 nm; Band 3: rest of the PAR region.

| C1B shade | Epidermis band 1 | Gastrodermis band1 | Epidermis band 2 & 3 | Gastrodermis band 3_2 | Assumptions |  |  |
| --- | --- | --- | --- | --- | --- | --- | --- |
| n | 1.38 | 1.42 | 1.38 | 1.42 | 1. Very low HP concentration.<br>2. Healthy symbiont density: 1X10<sup>6</sup> cells/cm<sup>2</sup> . | **Total source Intensity:**<br><u>1800 μmolm<sup>-2</sup>s<sup>-1</sup> / 400 Wm<sup>-2</sup></u><br>**Band 1: 120 Wm<sup>-2</sup>**<br>**Band2: 80 Wm<sup>-2</sup>**<br>**Band 3: 200 Wm<sup>-2</sup>** |  |
| μa (cm<sup>-1</sup>) | 0.0036 | 75 | 0.0036 | 54 |  |  |  |
| μs (cm<sup>-1</sup>) | 0.126 | 97 | 0.126 | 97 |  |  |  |
| g | 0.96 | 0.98 | 0.96 | 0.98 |  |  |  |
| C1B light | Epidermis band 1 | Gastrodermis band1 | Epidermis band 2 & 3 | Gastrodermis band 2 & 3 | Assumptions |  |  |
| n | 1.38 | 1.42 | 1.38 | 1.42 | 1. low HP concentration.<br>2. Partially bleached symbiont density: 0.5X10<sup>6</sup> cells/cm<sup>2</sup> . |  |  |
| μa (cm<sup>-1</sup>) | 0.0036 | 38 | 0.0036 | 26 |  |  |  |
| μs (cm<sup>-1</sup>) | 1.26 | 49 | 1.26 | 49 |  |  |  |
| g | 0.96 | 0.98 | 0.96 | 0.98 |  |  |  |
| C1G shade | Epidermis band 1_FL | Gastrodermis Band1_FL | Epidermis band 2_FL | Gastrodermis Band2_FL | Epidermis band 3_FL | Gastrodermis band 3_FL | Assumptions |
| n | 1.38 | 1.42 | 1.38 | 1.42 | 1.38 | 1.42 | 1. high FP concentration.<br>2. Healthy to partially bleached symbiont density: 0.75X10<sup>6</sup> cells/cm<sup>2</sup> . |
| μa (cm<sup>-1</sup>) | 0.0036 | 55 | 10 | 55 | 0.0036 | 40 |  |
| μs (cm<sup>-1</sup>) | 126 | 72 | 126 | 72 | 126 | 72 |  |
| g | 0.96 | 0.98 | 0.96 | 0.98 | 0.96 | 0.98 |  |
| C1G light | Epidermis band 1_FL | Gastrodermis Band1_FL | Epidermis band 2_FL | Gastrodermis Band2_FL | Epidermis band 3_FL | Gastrodermis band 3_FL | Assumptions |
| n | 1.38 | 1.42 | 1.38 | 1.42 | 1.38 | 1.42 | 1. high FP concentration.<br>2. Mostly bleached symbiont density: 0.25X10<sup>6</sup> cells/cm<sup>2</sup> . |
| μa (cm<sup>-1</sup>) | 0.0036 | 19 | 100 | 19 | 0.0036 | 11 |  |
| μs (cm<sup>-1</sup>) | 1260 | 24 | 1260 | 24 | 1260 | 24 |  |
| g | 0.96 | 0.98 | 0.96 | 0.98 | 0.96 | 0.98 |  |

Scalar irradiance measurements at the tissue-water interface further revealed FP-driven spectral redistribution (Fig. 4B). In the weakly-fluorescent CB morphs, relative scalar irradiance remained close to unity across the PAR region. In contrast, the fluorescent CG phenotype exhibited reduced scalar irradiance within the FP excitation and chlorophyll absorption ranges, and increased scalar irradiance within the FP emission range, reflecting fluorescence-mediated photon redistribution (but not unique evidence for that mechanism in these broadband measurements). Elevated FP concentrations also enhanced broadband light scattering, resulting in an overall increase in normalized scalar irradiance to approximately 1.5 at wavelengths outside the chlorophyll absorption band. Thus, FP expression primarily reshapes the spectral distribution of the internal light field rather than uniformly increasing or decreasing its magnitude.

These optical differences were reflected in the thermal response under high irradiance (1800 µmol photons m⁻² s⁻¹; Fig. 4A). In the CB phenotype, the shaded fragment exhibited higher temperature increases (∼1.0 ± 0.05 °C) than the light-exposed fragment (∼0.6 ± 0.05 °C), consistent with higher symbiont absorption. In contrast, the CG phenotype showed similar temperature increases under shaded and light-exposed conditions (∼0.67 ± 0.05 °C and ∼0.7 ± 0.1 °C, respectively), despite reduced symbiont density in the light-exposed fragment, a pattern consistent with, but not isolating, an additional FP-related contribution. We conclude that heat generation in weakly-fluorescent tissue is primarily governed by symbiont density, while high FP concentrations introduce a substantial, additional source of heating. However, these phenotype comparisons do not isolate pigment-specific heating: symbiont abundance covaries with light history, and the CB morph contains small amounts of cyan FPs that are not included in the model parameterization.

Interestingly, the highest temperature increase was observed in the shaded brown morph (CB_shade), which also exhibited the highest symbiont density (as estimated from reflectance spectra). It is therefore conceivable that the reduced symbiont densities observed in light-exposed colonies represent, at least in part, an acclimatory strategy to limit excessive heat generation. In this context, elevated FP expression may serve a complementary role by modifying the internal spectral distribution of light through wavelength-specific absorption and fluorescence emission, potentially reducing exposure to high-energy blue and UV wavelengths. However, at high concentrations, FPs may also contribute to increased tissue heating through the combined effects of enhanced scattering, absorption, and fluorescence-mediated redistribution of light. The net thermal response depends on the balance between light scattering, absorption, fluorescence emission, and symbiont abundance. This remains a hypothesis as our study did not manipulate symbiont density or measure bleaching, survival or thermal tolerance.

Simulations (Fig. 4C–F) reproduced the measured trends in surface scalar irradiance and temperature, providing mechanistic insight into the underlying processes. The optical and model parameters used for the simulation are shown in tables 1, 2 and 5. The defined wavebands represent key spectral regions: band 1 corresponds to the principal absorption range of symbiont pigments, band 2 to the absorption range of FPs and symbionts, and band 3 to the remaining PAR wavelengths (Table 1). In the CB model parameterization (Fig. 4C, D), light transport is governed primarily by scattering and symbiont absorption. Under shaded conditions, high symbiont density strongly attenuates scalar irradiance with tissue depth, leading to higher heat generation. Under light-exposed conditions, reduced symbiont density allows deeper light penetration but results in lower overall heat deposition due to weaker absorption.

In the fluorescent CG phenotype (Fig. 4E, F), FP absorption and fluorescence emission significantly modify both the magnitude and spectral distribution of scalar irradiance. Under shaded conditions (Fig. 4E, S2A), higher symbiont density dominates light attenuation, while the contribution of FPs remains limited because of their relatively low concentration. In contrast, the light-exposed CG fragment (Fig. 4F; CG_light_HF), characterized by reduced symbiont density, exhibits pronounced fluorescence-mediated spectral redistribution. Scalar irradiance within the FP excitation range (∼450–500 nm) is reduced near the tissue surface due to FP absorption, while fluorescence emission increases irradiance at longer wavelengths.

Hyperspectral and confocal imaging studies showed that FPs can occur either as diffuse tissue distributions or as localized granules, depending on the coral species and their spatial location within the tissue.^19,21^ Simulations were performed for both these scenarios. For the diffuse FP distribution assumed in the main model (Table 5), the local simulated irradiance in the fluorescence emission band reaches up to ∼2.5 times the incident excitation intensity within the FP absorption band (i.e., waveband 2). This ratio represents a comparison of local light fields rather than an energy-conversion fraction and does not imply that the emitted fluorescence energy exceeds the absorbed excitation energy. Rather, the elevated local fluorescence irradiance results from the redistribution and accumulation of emitted photons within the scattering tissue. In comparison, the granular FP configuration (Fig. S2B) produces fluorescence intensities of up to ∼7 times the incident excitation intensity for the same FP concentration. This stronger enhancement is attributed to increased backscattering and photon trapping within the upper epidermal layer, arising from the more isotropic scattering of granular FPs (g = 0.34), compared with the highly forward-scattering diffuse FP distribution (g = 0.96).

Fluorescence photons emitted following FP absorption can penetrate deeper into deeper tissue layers, where they are subsequently absorbed by symbionts, increasing subsurface energy deposition. Despite the increased backscattering associated with elevated FP concentrations, in the simulation scenarios, fluorescence-mediated spectral redistribution compensates for the reduction in symbiont light absorption caused by lower symbiont density.

Consequently, the simulated temperature profiles indicate that heat generation in CG_light_HF remains comparable to that in CG_shade, consistent with the experimental observations. The granular FP configuration produced slightly lower temperatures than the diffuse FP scenario due to increased photon backscattering and reduced net energy deposition within the tissue. Comparison of simulations with and without fluorescence re-emission (T_FL and T_noFL; where the latter is analogous to chromoproteins because all the absorbed energy in the epidermis is converted to heat like in CP model parametrization) further demonstrates that fluorescence contributes to tissue heating by redistributing absorbed and reemitted photons deeper into the tissue, where they are subsequently absorbed. As a result, tissue heating exceeds that arising from absorption alone, as in the corresponding chromoprotein scenario, with the magnitude of the effect depending on FP concentration. Thus, the model predicts a temperature difference between T_FL and T_noFL for these assumed FP parameters.

To further assess the role of FP concentration, additional simulations were performed for the light-exposed fluorescent phenotype (CG_light), while maintaining the same reduced symbiont density (Fig. 5, S2C–D). As FP concentration was not measured directly, it was varied as a model fitting parameter to reproduce the observed temperature profiles. Three FP-expression scenarios were considered: (i) low FP concentration (LF), corresponding to the concentration estimated for the shaded fluorescent phenotype (CG_shade), (ii) intermediate FP concentration (IF), representing a concentration midway between CG_shade and CG_light_HF, and (iii) high FP concentration (HF), corresponding to the concentration estimated for CG_light_HF and used in the simulations shown in Fig. 4F. This analysis allowed us to assess the sensitivity of fluorescence-mediated heating to FP abundance.

**Figure 5.**
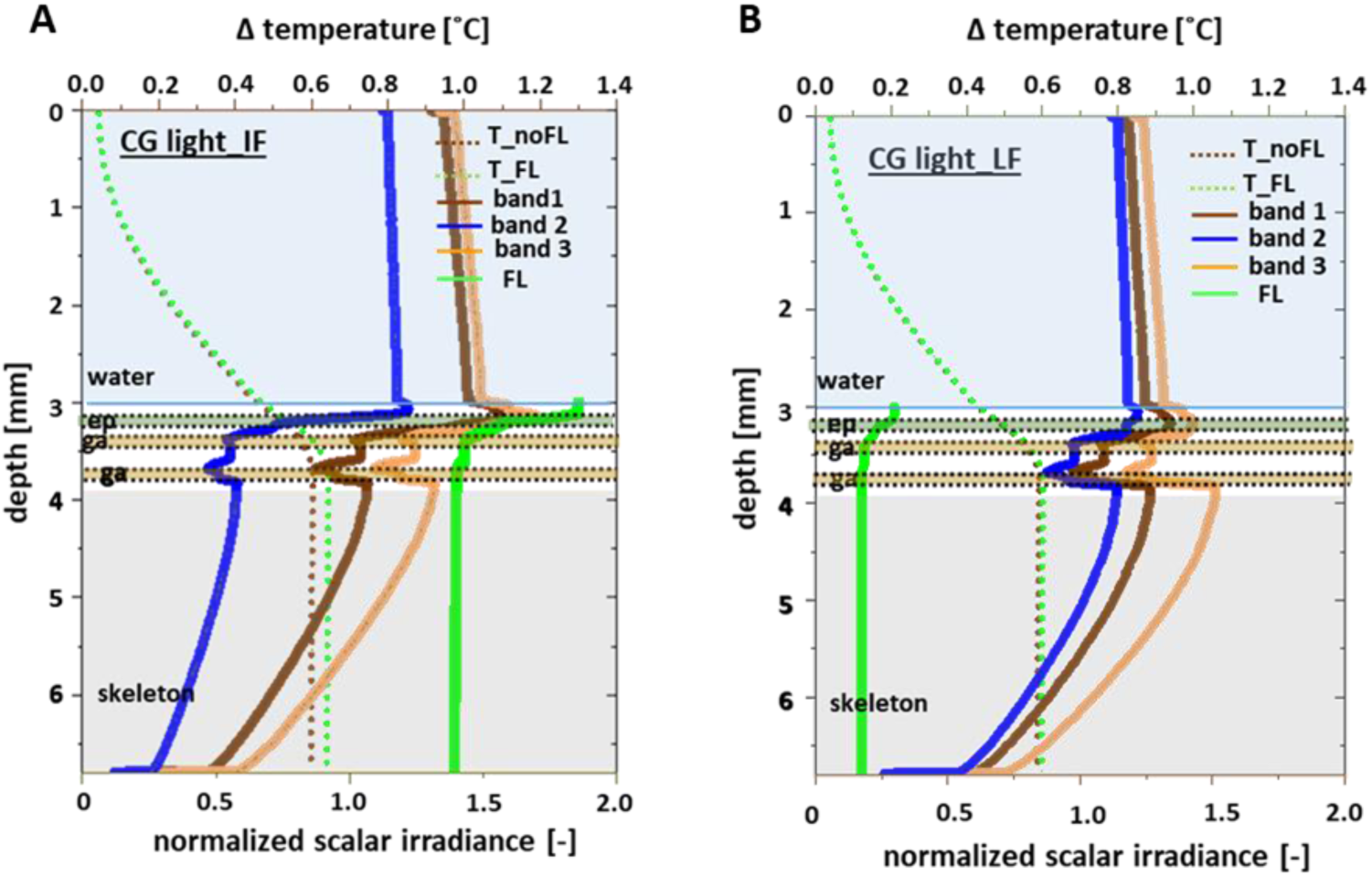
Scenario simulations examining the effects of different fluorescent protein (FP) concentrations in a diffuse FP configuration within a high light-exposed, fluorescent C. microphtalma (CG_light). (A) Simulated depth profiles of normalized scalar irradiance (bands 1–3 and fluorescence emission, FL) and temperature for CG_light fragment assuming a lower FP (IF) concentration, intermediate between that assumed for CG_shade and CG_light conditions shown in Fig. 4 (B) Simulated depth profiles of normalized scalar irradiance (bands 1–3 and fluorescence emission, FL) and temperature for CG_light assuming low FP concentration (LF, CG_shade concentration), while keeping the symbiont density identical to that of the measured CG light condition. Temperature profiles are shown for simulations including fluorescence re-emission (T_FL) and without fluorescence re-emission (T_noFL). Downwelling irradiance (1800 µmol photons m⁻² s⁻¹ / 400 W m⁻²). Band 1: 400–450 nm and 650–680 nm, principal absorption maxima of Symbiodiniaceae pigments; Band 2: 450–500 nm, absorption range for FP and Symbiodiniaceae pigments; band 3: remaining wavelengths in the PAR range (400-700 nm); ep: epidermis; ga: gastrodermis.

In the LF scenario (Fig. 5B, S2D), fluorescence-mediated spectral redistribution was weak due to the low FP concentration. Consequently, only a small fraction of incident photons was re-emitted at longer wavelengths, resulting in limited subsurface energy deposition and lower simulated heating. Although reduced symbiont density permitted deeper light penetration, the absence of substantial FP absorption and fluorescence prevented compensation for the reduced symbiont-driven energy absorption. The resulting temperature profile remained considerably lower than that measured experimentally for CG_light. In the IF scenario (Fig. 5A), moderate FP absorption within the excitation band increased fluorescence emission and photon redistribution to longer wavelengths. Compared with the LF case, this resulted in greater subsurface absorption and enhanced heat generation. The corresponding temperature profile was intermediate between the LF and HF scenarios, indicating a progressive increase in fluorescence-mediated heating with increasing FP concentration. In the HF scenario, corresponding to the FP concentration estimated for CG_light and used in Fig. 4F, strong fluorescence-mediated spectral redistribution was observed. Absorption of photons within the FP excitation range generated substantial fluorescence emission, increasing irradiance at longer wavelengths and promoting deeper photon penetration into the tissue. These re-emitted photons can subsequently be absorbed by the symbionts, enhancing subsurface energy deposition. The simulated surface temperature closely matched the experimentally observed heating of the light-exposed fluorescent phenotype despite its reduced symbiont density.

Together, these simulations demonstrate that FPs can influence the internal light and thermal microenvironment of corals in a concentration-dependent manner under the assumed optical conditions. At low FP concentrations, simulated tissue heating is primarily governed by symbiont abundance. As FP concentration increases, fluorescence-mediated spectral redistribution contributes progressively more to subsurface energy deposition and tissue heating. In the high-FP (CG_light_HF) scenario, the simulated temperature profile approaches that of the shaded coral with higher symbiont density (CG_shade), suggesting that elevated FP concentrations could partially compensate for reduced symbiont-mediated heating. However, because FP concentration was adjusted to reproduce the measured surface temperature, this agreement reflects model fitting rather than independent validation of the proposed mechanism. These simulations therefore illustrate the sensitivity of the thermal response to assumed FP concentrations rather than establishing an in vivo concentration-dependent relationship. We did not investigate potential saturation effects at higher FP concentrations, where increased scattering and reduced light penetration into deeper tissue layers could limit further increases in subsurface energy deposition and heating.^26^

#### 3.2.2 Astraeosmilia curvata

Individual *A. curvata* polyps exhibited two adjacent regions with comparable symbiont densities (inferred from reflectance measurements) but distinct host pigment composition; these were primarily FPs (Fig. 6). Although slight differences in symbiont density may exist, the optical measurements indicated that FP abundance is the dominant variable distinguishing the two regions. The green region (RG) showed bright green fluorescence, the brown region (RB) showed no detectable green fluorescence under the imaging conditions used (Fig. 6A, B).) Despite similar symbiont concentrations, the fluorescent region (RG) exhibited a slightly greater mean surface temperature increase (0.25 ± 0.03 °C) than the non-fluorescent region (RB; 0.20 ± 0.02 °C) under a downwelling irradiance of 600 µmol photons m⁻² s⁻¹. These values represent the mean ± SD of 12 measurements obtained from two coral fragments. Measurements were conducted in adjacent regions under comparable flow conditions. This trend is consistent with an additional contribution of FP expression to tissue heating beyond symbiont absorption.

**Figure 6.**
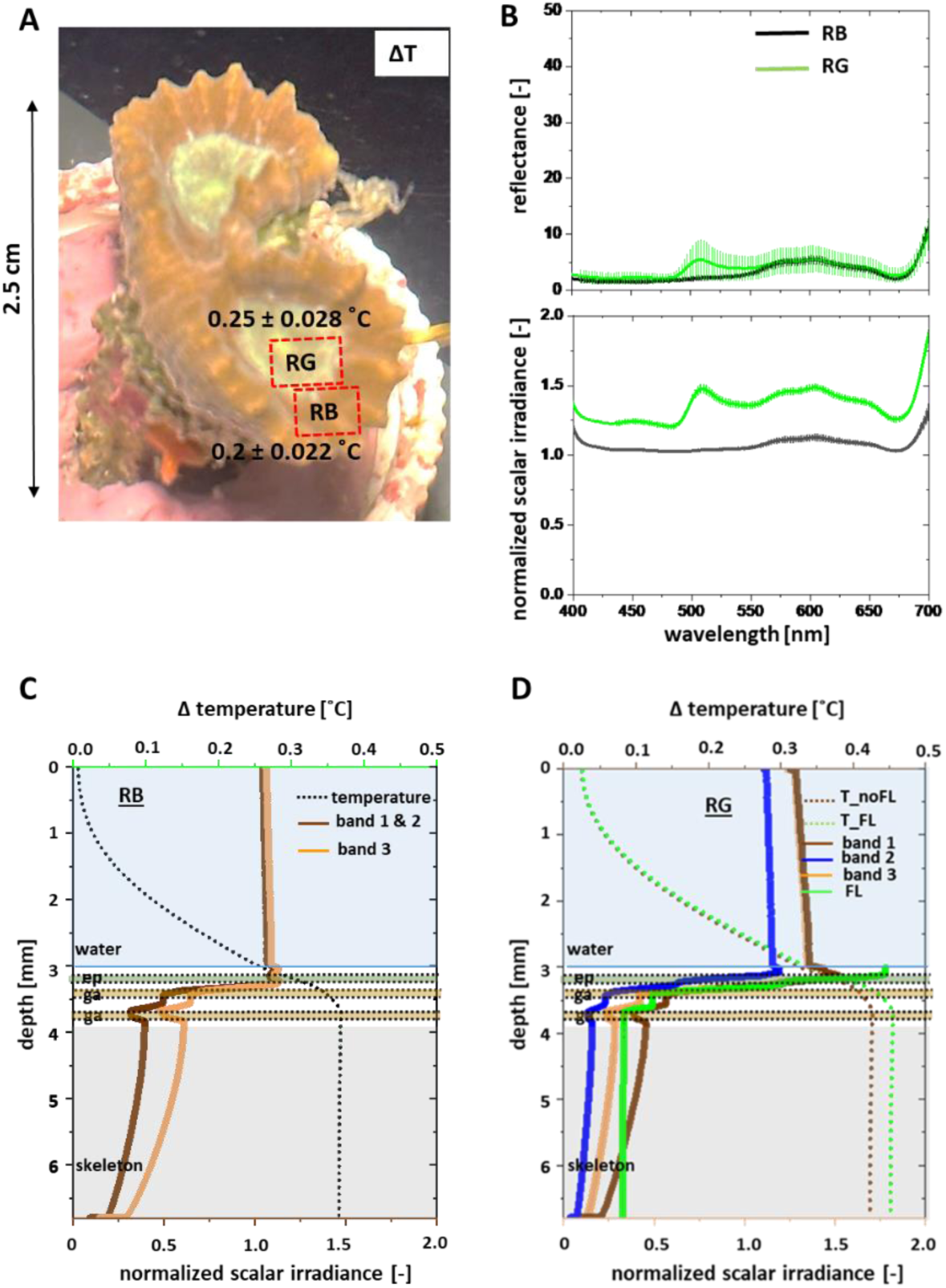
Optical and thermal effects of diffuse fluorescent proteins in Astraeosmilia curvata fragment (A) Representative image of a A. curvata fragment showing two distinct regions: a green, fluorescent region (RG) and a non-fluorescent, brown region (RB), with corresponding surface temperature increases (ΔT) under a downwelling irradiance of 600 µmol photons m⁻² s⁻¹ (130 W m⁻²). (B) Measured reflectance and normalized scalar irradiance spectra at the tissue-water interface for RG and RB. (C-D) Simulated depth profiles of scalar irradiance (bands 1–3) and temperature for RB and RG, respectively. Band 1: 400–450 nm and 650–680 nm, principal absorption maxima of Symbiodiniaceae pigments; Band 2: 450–500 nm, absorption range for FP and Symbiodiniaceae pigments; band 3: remaining wavelengths in the PAR range (400-700 nm); ep: epidermis; ga: gastrodermis. g = 0.96.

The reflectance measurements (Fig. 6B) illustrate these differences. Both regions showed strong attenuation of blue and red wavelengths associated with chlorophyll absorption, consistent with similar symbiont densities. However, RG exhibited distinct spectral features due to FP absorption in the excitation range (∼450–500 nm) and a pronounced emission peak at longer wavelengths (∼510–550 nm). Accordingly, scalar irradiance in RG was reduced within the excitation band and enhanced in the emission range, indicating fluorescence-mediated redistribution of photons within the tissue. The RG exhibited higher scalar irradiance across the PAR region than the RB region, owing to increased backscattering caused by the elevated concentration of FPs. In contrast, RB lacked this spectral redistribution, and its internal light field was governed primarily by scattering and symbiont absorption.

Computer simulations with our 2D model further elucidated these mechanisms (Fig. 6C, D). The optical and model parameters are provided in Tables 1, 2, and 6. The defined wavebands represent key spectral regions: band 1 corresponds to the principal absorption range of symbiont pigments, band 2 to the combined absorption range of FPs and symbionts, and band 3 to the remaining PAR wavelengths (Table 1). The simulated light and temperature profiles closely matched the experimentally measured scalar irradiance and surface temperature. Although the simulated surface temperatures were slightly higher than the measurements for both the RB and RG, this discrepancy is likely due to the symbiont densities assumed from the reflectance measurements. The aquarium-reared coral fragments bleached rapidly between measurements under the relatively high light intensity used. The temperature measurements were performed shortly after the reflectance and scalar irradiance measurements (all within 30 minutes), and during this time further changes in symbiont density may have occurred. The model reproduces the measured surface difference under its assigned optical parameters, but this is not independent validation. Nevertheless, the relative temperature differences between the RB and RG morphs are well reproduced by the simulations. In the non-fluorescent RB region (Fig. 6C), scalar irradiance decreases rapidly with depth due to strong symbiont absorption across the PAR spectrum, particularly in bands 1 and 2, leading to heat generation. In the fluorescent RG region (Fig. 6D), FP absorption reduces scalar irradiance near the surface within the excitation band, while fluorescence emission introduces additional irradiance at longer wavelengths. These re-emitted photons penetrate deeper into the tissue and are subsequently absorbed, enhancing energy deposition at depth.

**Table 6:**
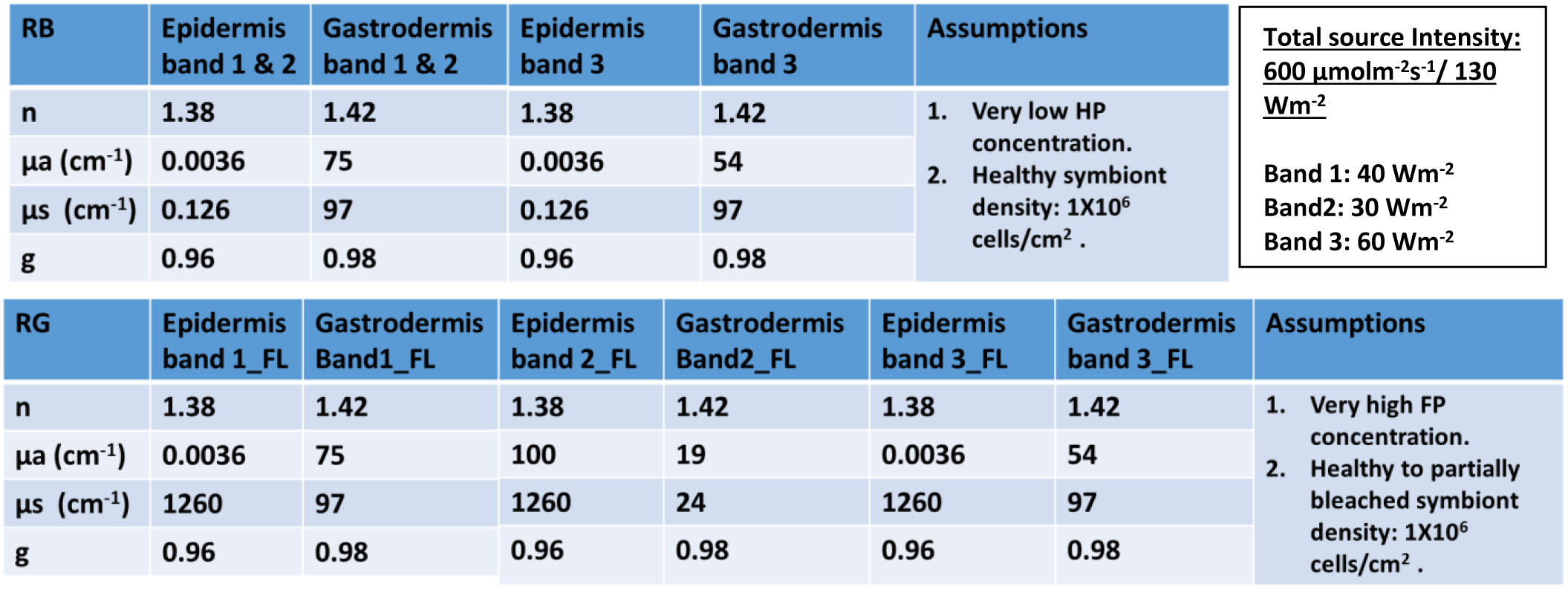
Summary of the epidermal and gastrodermal optical properties of the A. cuvata fragment, aquarium bred, with two distinct regions, with FP concentrations as indicated by reflectance spectra and visual inspection. RB: Region with healthy symbiont density and very low FP concentration. RG: Region with healthy symbiont density and very high FP concentration. Optical properties were determined as described in our previous work^1,2^. The epidermal absorption coefficient (µ_a_) was calculated as described in methods section.µ_s_: scattering coefficient; g: anisotropy of scattering and n: refractive index. The table also indicates the incident light intensity for each waveband in the PAR region. Band 1: 400 – 450 nm & 650 – 680 nm; Band 2: 450 – 500 nm; Band 3: rest of the PAR region.

| RB | Epidermis band 1 & 2 | Gastrodermis band 1 & 2 | Epidermis band 3 | Gastrodermis band 3 | Assumptions |  | Total source Intensity:<br>600 $\mu\text{molm}^{-2}\text{s}^{-1}$/ 130 $\text{Wm}^{-2}$<br>Band 1: 40 $\text{Wm}^{-2}$<br>Band2: 30 $\text{Wm}^{-2}$<br>Band 3: 60 $\text{Wm}^{-2}$ |
| --- | --- | --- | --- | --- | --- | --- | --- |
| n | 1.38 | 1.42 | 1.38 | 1.42 | 1. Very low HP concentration.<br>2. Healthy symbiont density: $1\text{X}10^6$ cells/$\text{cm}^2$ . |  |  |
| $\mu\text{a}$ ($\text{cm}^{-1}$) | 0.0036 | 75 | 0.0036 | 54 | | | |
| $\mu\text{s}$ ($\text{cm}^{-1}$) | 0.126 | 97 | 0.126 | 97 | | | |
| g | 0.96 | 0.98 | 0.96 | 0.98 |  |  |  |
| RG | Epidermis band 1_FL | Gastrodermis Band1_FL | Epidermis band 2_FL | Gastrodermis Band2_FL | Epidermis band 3_FL | Gastrodermis band 3_FL | Assumptions |
| n | 1.38 | 1.42 | 1.38 | 1.42 | 1.38 | 1.42 | 1. Very high FP concentration.<br>2. Healthy to partially bleached symbiont density: $1\text{X}10^6$ cells/$\text{cm}^2$ . |
| $\mu\text{a}$ ($\text{cm}^{-1}$) | 0.0036 | 75 | 100 | 19 | 0.0036 | 54 | |
| $\mu\text{s}$ ($\text{cm}^{-1}$) | 1260 | 97 | 1260 | 24 | 1260 | 97 | |
| g | 0.96 | 0.98 | 0.96 | 0.98 | 0.96 | 0.98 |  |

Importantly, comparison of the simulated temperature profiles with (T_FL) and without (T_noFL) fluorescence re-emission (the latter analogous to chromoproteins) demonstrates that, for the diffuse FP configuration (g = 0.96) fluorescence-mediated light redistribution enhances tissue heating in the fluorescent RG region beyond that produced by absorption alone, with the magnitude of the effect depending on FP concentration. For the granular FP configuration (g = 0.34; Fig. S3), the simulated RG temperatures with and without fluorescence re-emission are comparable to those of the RB region because the more isotropic scattering by FP granules increases photon backscattering, reducing light penetration into the tissue. Nevertheless, the simulated temperature with fluorescence (T_FL) remained slightly higher than that without fluorescence (T_noFL), indicating the contribution of fluorescence-mediated light redistribution. Overall, the predicted temperatures were lower than those for the diffuse FP configuration owing to the greater loss of optical energy through backscattering.

Together, these results demonstrate that FP expression can modify the internal light field independently of major differences in symbiont abundance. By absorbing photons within the excitation range and re-emitting a fraction of the absorbed energy as fluorescence, FPs alter both the spectral and spatial distribution of light within the tissue. The resulting thermal response reflects a balance between broadband light scattering, wavelength-specific absorption, and fluorescence-mediated photon redistribution. While enhanced scattering increases photon escape from the tissue, fluorescence redistributes a portion of the absorbed energy to deeper tissue layers, where it can be subsequently absorbed by the symbionts, increasing subsurface energy deposition and tissue heating. Consequently, the net optical and thermal effects of FPs depend on their concentration, with higher FP concentrations producing a greater fluorescence-mediated contribution to tissue heating. Unlike chromoproteins, which generate heat primarily within the superficial epidermis where it can be readily dissipated to the surrounding seawater, FPs re-emit a fraction of the absorbed energy as fluorescence. These fluorescence photons penetrate deeper into the tissue, where their subsequent absorption increases subsurface energy deposition and tissue heating.

## 4. Conclusions and outlook

This study demonstrates that synthesis of host pigments, i.e., chromoproteins (CPs) and fluorescent proteins (FPs), enables different optical mechanisms for regulating the internal light and thermal microenvironment of corals. By combining microscale optical and thermal measurements with mechanistic light transport and heat transfer modelling, we show that tissue heating is governed by the interplay between broadband scattering, wavelength-specific absorption by host pigments and symbionts, and, in the case of FPs, fluorescence-mediated photon redistribution.

While CP absorption generates localized heat primarily within the superficial epidermis, symbiont absorption produces heat deeper within the gastrodermis, where thermal dissipation is less efficient. Unlike CPs, FPs additionally redistribute a fraction of the absorbed energy through fluorescence, shifting energy deposition deeper into the tissue and enhancing heating relative to the corresponding non-fluorescent (CP) scenario. The magnitude of this enhancement depends on FP concentration and spatial organization, reflecting the trade-off between fluorescence-mediated photon redistribution and photon loss through elastic backscattering. Together, these findings demonstrate that coral host pigments can regulate not only the amount of absorbed light but also where optical energy is deposited within the tissue, providing a mechanistic framework for understanding coral photoacclimation and bleaching resilience.

These findings establish a quantitative link between host pigment composition, internal light availability, and tissue heating within coral tissues, with implications for coral performance under environmental stress, consistent with previous studies demonstrating strong optical feedbacks in coral tissues associated with bleaching and host pigment upregulation.^3,11,28,29,32^ By resolving depth-dependent light redistribution and heat deposition (simulated, only surface measurements were made), the results show that pigment-driven optical feedbacks can either dampen or sustain internal energy loads depending on pigment type, concentration, and symbiont density. Such effects are likely to influence thermal stress responses, photosynthetic light availability, photosynthetic performance, and recovery following bleaching.^22,41^ This study does not measure photosynthesis, thermal tolerance, or recovery; it therefore cannot establish whether the predicted redistribution improves or reduces bleaching resilience.

Future work should extend this modeling framework to more complex and realistic coral architectures.^12^ Incorporating fully three-dimensional geometries derived from high-resolution scans will enable investigation of how skeletal structure, tissue microtopography, and polyp-scale morphology influence light redistribution and localized heat generation.^7,12,42^ In addition, integrating other biologically relevant components, such as endolithic algae^43,44^ and dynamic tissue optics^40,45^, will further improve model realism and predictive capability.

Beyond radiative and thermal processes, an important next step is to couple host pigment modulated light transport with physiological processes occurring within the tissue^19,46^ and bleaching response^21,28^. Changes in the internal light field directly influence photosynthesis, oxygen production, and pH dynamics, all of which affect symbiont performance and calcification.^6,33,47^ Incorporating these processes into the modeling framework would enable assessment of how host pigment driven modifications of the optical microenvironment influence carbon fixation, oxygen availability, and the chemical conditions experienced by both the coral host and its symbionts under changing environmental conditions.^48^

Finally, the framework developed here offers a foundation for predictive, “digital twin”-type approaches to coral function, integrating optical, thermal, and physiological processes across scales. It could support the design of targeted experiments to investigate pigment-mediated photoprotection and internal light and temperature conditions. However, validation against independent internal measurements and incorporation of a modeled bleaching or recovery response would be needed before using the framework to identify pigment-mediated pathways that improve thermal tolerance or to guide reef restoration strategies.

## 2. Materials and Methods

The methodological framework comprised both experimental and theoretical components. Microsensor measurements of light and temperature were first conducted on different coral fragments placed in a flow chamber under controlled irradiance and flow conditions (Figure S4). The same fragment was subsequently 3D scanned to obtain its geometric properties. A 2D slab model representing layered coral tissue overlying the skeleton was used to simulate radiative and heat transfer within the coral (Figure S5). The resulting spatial distributions of light and temperature were then quantitatively compared with corresponding experimental measurements.

### 2.1 Coral specimens

Corals adapted to high-light conditions were collected from the shallow reef-flat (< 2 m depth) at Heron Island, Great Barrier Reef, Australia (152°06′ E, 20°29′ S. Permits n.G24/49877.1; G26/52406.1). Under clear-sky conditions at midday, downwelling photon irradiance at this site can reach approximately 2000 μmol photons m⁻² s⁻¹, with light distributed relatively evenly across the photosynthetically active radiation (PAR) range (400–700 nm).^49^ To obtain representative samples of corals with different symbiont and host pigment concentrations, we collected fragments from multiple species of corals spanning healthy and bleached colonies, different colour morphs, and high light exposed as well as shaded regions of the colonies. The selected species comprised: a fragment of *Isopora palifera*, displaying a colourful bleaching phenotype on the light-exposed side and a brown phenotype on the underside; two fragments of a *Montipora capricornis* colony, one displaying a pink color from the light exposed region and the other a brown color from the shaded region; fragments of two color morphs (green and brown) of *Cyphastrea microphthalma*, collected from both the light exposed part and the shaded part of each colony. Prior to measurements, all coral fragments were transferred to a flow-through holding tank supplied with seawater from the reef and maintained under natural sunlight filtered through shade cloth to reach a midday photon irradiance of 350 µmol photons m⁻² s⁻¹. An aquarium-reared coral fragment of *Astraeosmilia cuvata* (discussed in example 4) was obtained from Dejong Marine Life (Netherlands).

### 2.2 Experimental setup

Fragments of different species of corals were placed, one at a time, in a custom-made black acrylic flow chamber (Figure S4A). Care was taken to prepare the coral fragments as flat specimens whenever possible. If fragmentation did not produce a sufficiently planar surface, the fragment was positioned flat within the flow chamber to minimize geometry-dependent variations in the local flow field and ensure comparable hydrodynamic conditions between samples. Maintaining similar flow conditions was essential to enable accurate comparisons between the colorfully bleached samples and their corresponding high symbiont density controls. During measurements, the coral was continuously flushed with aerated seawater collected from the reef. An average flow velocity of ∼1 mm s⁻¹ was sustained using a water pump. The fragment was allowed to acclimate in the chamber for at least 1 h prior to measurements to ensure steady-state conditions.

Illumination was provided by a fibre-optic tungsten-halogen lamp equipped with a heat filter and collimating lens (KL-2500 LCD, Schott GmbH, Germany), positioned vertically above the chamber (Figure S4A). The coral fragment was exposed to a constant downwelling photon irradiance of 600 µmol photons m⁻² s⁻¹ (aquarium sample) or 1600/1800 µmol photons m⁻² s⁻¹ (field samples). For each comparison, the illumination level was kept identical between the color morphs and its corresponding control. During all measurements, the region of interest was positioned centrally within the light beam to ensure uniform illumination. Under natural reef conditions, incident light typically fluctuates between collimated and diffuse fields and only reaches comparable irradiance levels on shallow reef flats at midday under clear skies and calm waters^14,50^. The relatively high and collimated irradiance applied here was selected to simplify boundary conditions and to induce measurable temperature elevations.

The experimental photon irradiance levels were obtained by adjusting the aperture of the fibre-optic halogen lamp, thereby modifying light intensity without altering the spectral distribution. Downwelling spectral irradiance at these photon flux densities was additionally quantified in radiometric units (W m⁻² nm⁻¹) using a calibrated spectroradiometer (360 - 830 nm, BTS256-PAR Meter, Gigahertz-Optik).

### 2.3 Microsensor measurements

Measurements were conducted on the coral tissue (coenosarc) between individual polyps. This tissue region was chosen because it had a relatively uniform surface topography and was less contractile than polyp tissue, thereby reducing variability during microsensor positioning and measurements.

Microsensors were mounted on a manually operated micromanipulator (MU-1, PyroScience GmbH, Germany). During measurements, the sensors were positioned at a 45° angle relative to the vertically incident light beam to minimize self-shading.

Scalar irradiance, reflectance, and temperature were measured by carefully positioning the sensor tip at the coral tissue surface. The position of the microsensor tip was continuously monitored under a dissection microscope to ensure precise placement. At least six measurements were taken at different positions across each sample (n ≥ 6) and averaged.

### 2.4 Reflectance measurements and optical properties

Reflectance was measured using optical fiber– based spectroscopy. A flat-tip fiber optic field radiance probe^51^ (1 mm diameter) was connected to a spectrometer (USB 2000+, Ocean Optics, USA). The incident light spectrum was recorded at the same height as the coral surface by placing the sensor above a 99% reflectance standard (Spectralon, Labsphere) in air. All measurements were dark-corrected and normalized to the corresponding dark-corrected incident spectrum. Reflectance values within the photosynthetically active radiation (PAR; 400–700 nm) range are reported as mean ± standard deviation.

Reflectance spectra (which also includes fluorescence contribution by HPs) were used to guide assumptions about the optical properties of the tissue sublayers used in the numerical modeling (see section 2.7), i.e., the absorption coefficient (µ_a_) and scattering coefficient (µ_s_). Relative changes in host pigment concentration within the epidermis and symbiont density within the gastrodermis were estimated from reflectance measurements at their respective absorption wavelengths. These estimates were then used to scale the optical properties assigned to the model layers. The resulting values should therefore be interpreted as comparative, relative changes between the morphotypes rather than absolute concentrations or optical property measurements. Chromoproteins (CPs) examined here have an absorption band 540–620 nm^28^, while fluorescent proteins (FPs) have an absorption band 450–500 nm^20,23^. The absorption features of symbiont cells are linked to chlorophyll a, chlorophyll c, and carotenoids, primarily in the ranges 400–490 nm and 650–690 nm.^41^ Based on these fragment-specific assumptions, optical properties were assigned using literature values and implemented in Monte Carlo simulations of light transport.

To reduce computational cost while capturing the key factors governing the optical properties of the coral fragments, and to ensure comparability across samples, the PAR region (400-700 nm) was divided into three spectral bands.

For coral fragments exhibiting significant CP expression and their controls

i. Band 1_cp (400–490 nm and 650–680 nm): corresponding to the main absorption maxima of Symbiodiniaceae pigments^41^ (chlorophylls *a* and *c*; carotenoids);
ii. Band 2_cp (540–620 nm): representing the CP absorption range^28^;
iii. Band 3_cp: the remaining portion of the PAR spectrum not included in Bands 1 and 2.

For colorfully bleached fragments with elevated CP concentrations, the epidermal absorption coefficient (µ_a_) was calculated using Equation (1).^4^ Since these fragments were mostly or completely bleached with high expression of host proteins, the calculation assumed light absorption and scattering to occur in the epidermal layer, while the contribution from skeletal scattering is comparatively low (see optical properties in Table 3 and 4).

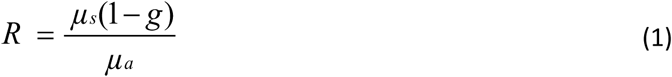

where *R* is the average reflectance (%) within the CP absorption band (540–620 nm)^28^ obtained from the reflectance spectra; µ_s_ is the scattering coefficient, assumed from literature values for highly upregulated CP conditions^1,52^, g= 0.96 assuming diffuse distribution of CPs^1,52^; µ_a_ is the absorption co-efficient of the epidermis layer.

For coral fragments exhibiting significant FP expression and their controls

i. Band 1_fp (400–450 nm and 650–680 nm), corresponding to the primary absorption maxima of Symbiodiniaceae pigments^41^ (chlorophylls *a* and *c*; carotenoids);
ii. Band 2_fp (450–500 nm), representing the FP absorption range^20,23^ and overlapping with absorption maxima of some of the Symbiodiniaceae pigments^41^;
iii. Band 3_fp, comprising the remaining portion of the PAR region not included in Bands 1 and 2.

For fragments with upregulated FP concentrations, the epidermal absorption coefficient (µ_a_) for band 2_fp was calculated using Equation 2^53^.

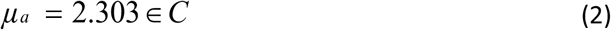

Where µ_a_ is the absorption co-efficient of the epidermis layer for band 2, ∈ is the molar extinction extinction coefficient, (approximated to 50,000 M⁻¹ cm⁻¹ for coral FPs^54^), and C is the FP concentration in the tissue. Since absolute FP concentrations in corals are not well reported, values of 100 µmol L⁻¹ and 1000 µmol L⁻¹ were assumed to represent high (normal, shaded) and very high (light-exposed) concentrations, respectively. These concentrations were treated as model parameters and adjusted to achieve the best agreement between the simulated and experimentally measured surface temperatures.^19,26^ Accordingly, µ_a_ values of 11 cm⁻¹ for high (shaded) and 110 cm⁻¹ for very high (light-exposed) concentrations were assumed.

The absorption coefficients of CPs and FPs were estimated using different approaches because their absorption bands differ in the extent to which they overlap with symbiont pigments. For CPs, absorption occurs primarily within the 540–620 nm spectral region, where symbiont absorption is relatively weak. Consequently, in the predominantly bleached, CP-rich fragments examined here, CP absorption could be estimated directly from the reflectance spectra using equation 1. In contrast, the excitation band of FPs overlaps substantially with major symbiont absorption bands, and the fluorescent phenotypes considered in this study retained sufficient symbiont densities that their contribution could not be neglected. Therefore, FP absorption could not be directly inferred from reflectance measurements alone and was instead estimated using equation 2 based on relative FP concentrations and extinction coefficients from literature.

For both CPs and FPs, the resulting optical properties should be interpreted as effective microscale model parameters describing relative differences among phenotypes rather than absolute measurements of pigment concentration or tissue optical properties. Agreement with surface scalar irradiance and temperature is a consistency check, not independent validation, because some optical parameters were inferred or fitted using these measurements; internal depth profiles were not measured.

The wavelength specific optical properties (µ_a_, µ_s,_ g and n) for high symbiont density coral fragments with low HP concentrations are listed in Table 1. Fragment-specific optical properties across different spectral bands for the epidermis and gastrodermis are presented in Tables 3–6. The optical properties of the remaining tissue sublayers and the skeleton were assumed to be identical across all fragments (Table 1).

Optical properties for varying host pigment and symbiont concentrations were adapted from our previous work.^1^

### 2.5 Scalar irradiance measurements and calculations

Spectral scalar irradiance was measured using a fiber-optic scalar irradiance microprobe with a spherical tip^55^ (∼100 µm diameter) connected to a fibre-optic spectrometer (USB2000+, Ocean Optics, USA). Incident downwelling irradiance was quantified at the same vertical position as the coral surface by placing the microsensor in a black, non-reflective light well beneath the vertically incident illumination within the flow chamber; the downwelling irradiance and downwelling scalar are identical in a collimated light field^56^. Scalar irradiance spectra recorded at the coral tissue surface were normalized to the corresponding incident spectral irradiance. Data are reported as mean ± standard deviation based on at least six measurement points within each defined region of interest for each fragment.

To express scalar irradiance in energy units, the recorded spectral counts at each wavelength were converted to absolute spectral scalar irradiance by multiplying normalized scalar irradiance spectra with calibrated spectral measurements of downwelling irradiance (W m⁻² nm⁻¹) obtained at the same vertical position using a calibrated spectroradiometer (BTS256-PAR Meter, Gigahertz-Optik). The total scalar irradiance (W m⁻²) within specific spectral regions was calculated by integrating the scalar irradiance spectra over defined wavebands within the PAR range. Integrated energy values were determined separately for each band, and the corresponding values for different coral fragments are presented in Tables 3–6.

### 2.6 Temperature measurements

Point measurements of temperature at the coral surface were carried out using thermocouple microsensors (tip diameter ∼200 µm; TP200, Unisense A/S) connected to a thermocouple meter (Unisense A/S). Temperature (ΔT) was defined as the increase relative to the water temperature in the free-flow region directly above the measurement point. At least five measurements were obtained for each region of interest on the coral fragments and averaged.

### 2.7 Numerical modeling

A two-dimensional model was developed to simulate the spatial distribution of light and temperature within and around the stratified coral tissue and skeleton. Model predictions were compared with measured surface scalar irradiance and temperature. This approach allowed us to assess how colorful bleaching modifies the surface and internal light and thermal microenvironment of the coral.

#### 2.7.1 Model geometry

The model geometry represents a coral slab composed of sequential layers of water, coral tissue, and skeleton (Figure S5A). As the coral fragments were mostly planar or positioned flat in the flow chamber during measurements, the effects of three-dimensional morphology on light distribution and flow were assumed to be negligible or comparable across fragments. Furthermore, at the considered spatial scale (milli-to micrometers), the tissue layers appear quasi-planar, therefore, a two-dimensional geometry was adopted for computational efficiency.

Water was defined as the only flowing domain. The coral tissue was represented as a stratified structure extending from the surrounding water to the skeleton surface, including the following functional layers: mucus, epidermis (containing fluorescent or non-fluorescent host pigments), oral mesoglea, gastrodermis with photosynthetic symbionts, a water-filled gastrovascular cavity, aboral gastrodermis (also containing symbionts), and aboral mesoglea (Figure S5A,B).^1,11,12^ The calicoblast was merged with the skeleton as its optical signature was assumed to derive mostly from aragonite. Most corals examined in this study, except *A. curvata*, are thin-tissued; therefore, the total tissue thickness was assumed to be ∼800 µm.^12^ The thicknesses of the individual sublayers are shown in Figure S5A. Although *A. cuvata* is typically a thick-tissue coral, the high light intensities used during measurements likely induced tissue contraction.^11,40^ Consequently, a thin-tissue model geometry was also applied in this case. The width of the 2D slab was chosen large enough to establish the thermal boundary layer. Host pigments were assigned only to the epidermis in this model; their precise location in these fragments was not established here. If pigments also occur in the gastrodermis, the model does not represent their local absorption, emission, or heat deposition.

#### 2.7.2 Model solution

A schematic of the slab model, including boundary conditions for light, flow, and heat transfer, is shown in Figure S5. A mesh-independence analysis for the base case indicated that a minimum element size of 2 µm was sufficient for Monte Carlo light simulations, while a minimum element size of 10 µm was used for flow and heat transfer simulations.^1^ All model parameters are listed in Tables 1 – 6.

##### Radiative transfer

Radiative transfer of incident light was simulated using a Monte Carlo (MC) approach (Figure S5B, C), as implemented in the open-source software ValoMC.^1,12,35^ Photon transport was modeled across the PAR range (400–700 nm), with optical properties for the different tissue layers and the skeleton (Tables 1–6) assigned to specific spectral bands as described in Section 2.4. The simulations produced spatial distributions of scalar irradiance within the coral and the surrounding water. The resulting point cloud of scalar irradiance values from ValoMC was imported into COMSOL (v 6.3; COMSOL Inc.) and mapped onto the 2D slab geometry using linear interpolation. Simulated scalar irradiance at each position was then normalized to the incident irradiance.

For fluorescence light transport simulations, coral tissue and skeleton were treated as highly scattering media, and fluorescence propagation was approximated using the diffusion equation derived from the radiative transfer equation (RTE).^34,57^ This approximation was applied to fluorescence emitted by green fluorescent proteins (FPs) in band 3_fp (emission peak ∼520 nm), where absorption is expected to be relatively weak compared with the principal absorption bands of FPs and *Symbiodiniaceae*. However, absorption within this spectral region is not negligible in the gastrodermis layer, as reflected by the nonzero gastrodermal absorption coefficients assigned in Table 1, and varies with symbiont concentration and pigment composition. However, scattering dominates absorption (µ_s_> µ_a_), allowing the radiative transfer equation (RTE) to be approximated by the diffusion equation.^34,57^ This approach is commonly employed in tissue optics for modeling light propagation in biological media.^34,57–59^

The simulated incident light field within the epidermis layer (I_ep_) was used to calculate the fluorescence emission, which was implemented as a secondary light source in the model (Figure S5 C,D), as described in Equation (3):

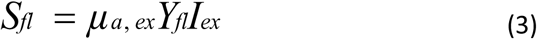

where S_fl_ is fluorescence source intensity, µ_a,ex_ is the absorption coefficient of the epidermis in band 2_fp (excitation wavelength,450 – 500 nm; calculated using equation 2), and Y_fl_ is the quantum yield of coral fluorescent proteins (0.65)^38^; I_ex_ represents the simulated normalized scalar irradiance (I_ep_) multiplied by the incident power of the excitation source in band 2_fp.

The fluorescence quantum yield was assumed to be 0.65 based on published values for green fluorescent proteins (FPs) from other cnidarian species.^38^ This value was used as a model parameter and was not experimentally determined for the FPs expressed in the coral fragments investigated here. Although Ilagan et al. (2010)^60^ reported varying quantum yields for different fluorescent proteins, including a value approaching 1.0 for an FP isolated from *Cyphastrea microphthalma*, the identity and in vivo quantum yield of the FPs in our samples remain unknown. Consequently, the simulated fluorescence-mediated light redistribution and associated tissue heating are conditional on the assumed quantum yield. Furthermore, because FP concentration was varied to match the measured surface temperatures, the fitted concentrations may depend on the assumed quantum yield and should not be interpreted as independently determined FP abundances.

Fluorophores in the epidermis were assumed to emit isotropically. The diffusion equation was solved in COMSOL Multiphysics using the Helmholtz formulation, equation 4:

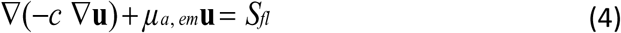

Where µ_a,em_ the absorption coefficient, of the respective layer, at the emission wavelengths (i.e. band3_fp), S_fl_ is the fluorescence source term, and *c* is the diffusion coefficient described in equation (5):

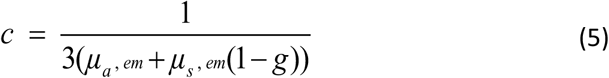

Where µ_a,em_ and µ_s,em_ are the absorption and scattering coefficients at the emission wavelength (band3_fp), for the respective layer, and g is the scattering anisotropy factor.

##### Fluid flow

The water flow around the coral fragment was simulated in COMSOL using the stationary incompressible Navier–Stokes equations (6 & 7) for laminar flow (Reynolds number ∼ 10).

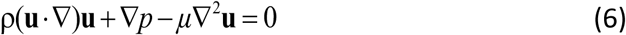

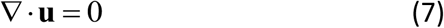

where u is the velocity vector, *p* is pressure, *μ* is the dynamic viscosity, and ρ is the density of water. A uniform inlet velocity of 1 mm s⁻¹ was applied, while the outlet was set to zero-gauge pressure. The top, bottom, and lateral walls of the flow cell were defined as no-slip boundaries (zero velocity).

##### Heat generation and transport

The spatial temperature distribution (T) within the coral and the surrounding water was computed using the general heat balance equation, accounting for conduction, convection, and internal heat source terms. In the coral tissue layers and skeleton (i.e., solids), the heat balance included heat conduction and heat generation terms (Eq. 8), whereas in the water layer (i.e., liquid), both heat conduction and convective heat transport were considered (Eq. 9).

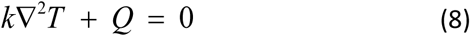

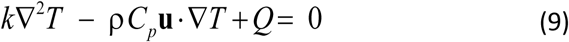

where *k* is the thermal conductivity of the respective layer, C_p_ is the specific heat capacity of water, Q is the heat source term, and u is the water velocity vector. The heat source was assumed to originate solely from light absorbed by the coral tissue and skeleton and was defined as

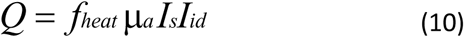

where μ_a_ is the absorption coefficient, *I_s_* is the normalized scalar irradiance, and *I_id_* is the incident downwelling irradiance (source). The fraction of absorbed light dissipated as heat (*f_heat_*) was set to 0.99 in the oral and aboral gastrodermis^15^ and to 1 in all other layers. An exception was made for the epidermis in the FP excitation band (band_fp) in samples with higher FP concentration, where f_heat_, representing the non-radiative quenching factor, was set to (1 − Y_fl_). Heat produced by biological metabolism was considered negligible^16^ (<1% of the total heat source). A value of 0.99 was assumed for the gastrodermis to account for the fraction of absorbed energy fixed through photosynthesis, as under high-light conditions, most of the absorbed energy is dissipated as heat. ^15^

The inflow of water had a temperature of *T_0_* (25 °C), while a zero-temperature-gradient condition was set for the water outflow. Thermal insulation was set at all other flow-cell walls, while heat flux continuity was assumed at the coral/water interface. The thermal properties are tabulated in Table 2

### 2.8 Comparison between simulated and measured data

For comparison between measurements and simulations, scalar irradiance was measured at the coral tissue-water interface at different wavelengths and normalized to the incident irradiance. These measurements were averaged (n ≥ 5) across defined regions of interest and compared with simulated scalar irradiance values extracted from the corresponding regions at the center of the 2D coral slab model.

For temperature, averaged point measurements (n ≥ 5) at the tissue-water interface over the same regions of interest were compared with simulated temperatures at the coral-water interface in the center of the slab model. In addition, depth profiles of scalar irradiance and temperature were extracted along the central axis of the 2D slab geometry to assess the vertical distribution of light and heat within the tissue and skeleton.

It should be noted that certain factors, such as the presence of endolithic algae, are not included in the model. Therefore, a direct one-to-one validation between simulations and measurements is neither feasible nor intended. Instead, the model is used to provide mechanistic insight into the colorful bleaching phenomenon and to clarify the roles of host pigments and symbionts in shaping internal optical properties and heat generation in corals.

### 2.9 3D texture scanning

A PC-controlled, structured light 3D scanner (EinScan-SP, Shining 3D) was used to generate high-resolution 3D models of coral fragments in air, with a spatial resolution of ∼50 µm. Each sample was placed on a calibrated turntable, and data acquisition and processing were performed using the system software (EXScan S_V3, Shining 3D). Scanner brightness was adjusted to avoid saturation of the camera signal. A full 360° scan of each fragment was obtained from multiple orientations to capture the complete morphology and minimize missing surfaces. Individual scans were aligned using the “align by feature” function in the software. In addition to geometry, texture scans were acquired using sequential red, green, and blue (RGB) structured illumination to capture surface colour information. These RGB projections enabled reconstruction of textured 3D models, preserving spatial variations in pigmentation across the coral surface. The combined geometry and texture data were processed to generate a watertight mesh, which was exported as OBJ and STL files.

## Acknowledgements

The study was supported by research grants from the Villum Foundation [VIL50371 & VIL57413; MK], and the Gordon and Betty Moore Foundation [grant no. GBMF9206; https://doi.org/10.37807/GBMF9206; MK]. We acknowledge the excellent assistance of the technical staff at Heron Island Research Station during field work, and the technical assistance of Sofie Jakobsen with coral husbandry and lab organization. The field work on Heron Island was conducted under permit no. G24/49877.1 and G26/52406.1 from the Great Barrier Reef Marine Parks authorities. The authors thank Rob Brennan for helping with sample collection.

## Author Contributions

SM, EB, AS and MK conceptualized the study. Samples were collected by SM, EB, AS, and MK. SM and EB optimized the measurement protocol. Measurements were performed by SM with input from EB. SM conducted data analysis and numerical simulations. All authors (SM, EB, AS, and MK) contributed to data interpretation and discussion. SM wrote the manuscript, with editorial input from EB, AS, and MK.

## Supplementary information

**Figure S1:**
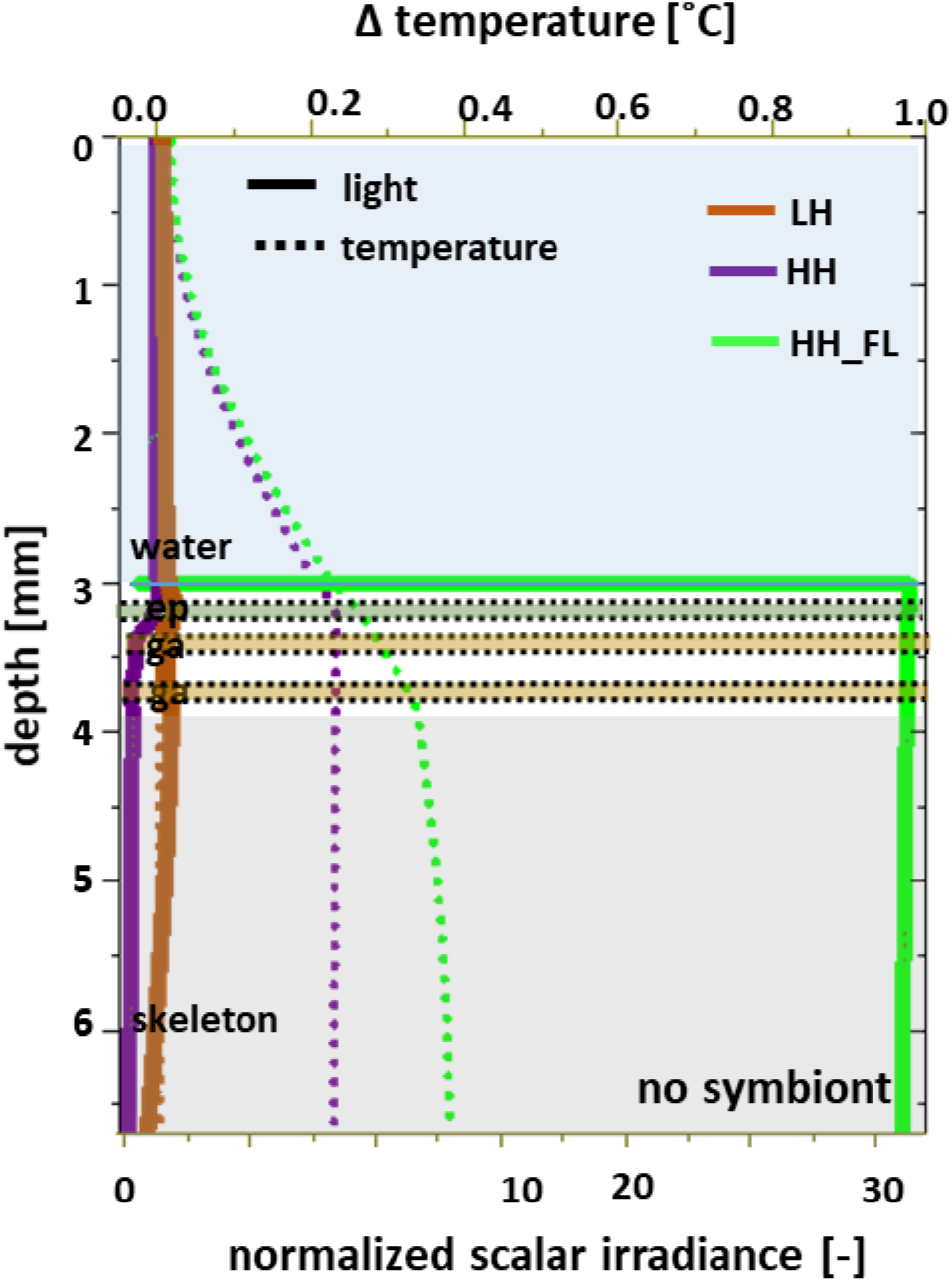
Simulated depth profiles of normalized scalar irradiance (PAR) and temperature in coral with no symbionts, comparing conditions with high HP concentrations with and without fluorescence, and with low HP concentrations. LH: fragment with low host protein concentration; HH: fragment with high host protein concentration, with no HP fluorescence; HH_FL: fragment with high host protein concentration, with HP fluorescence.

**Figure S2.**
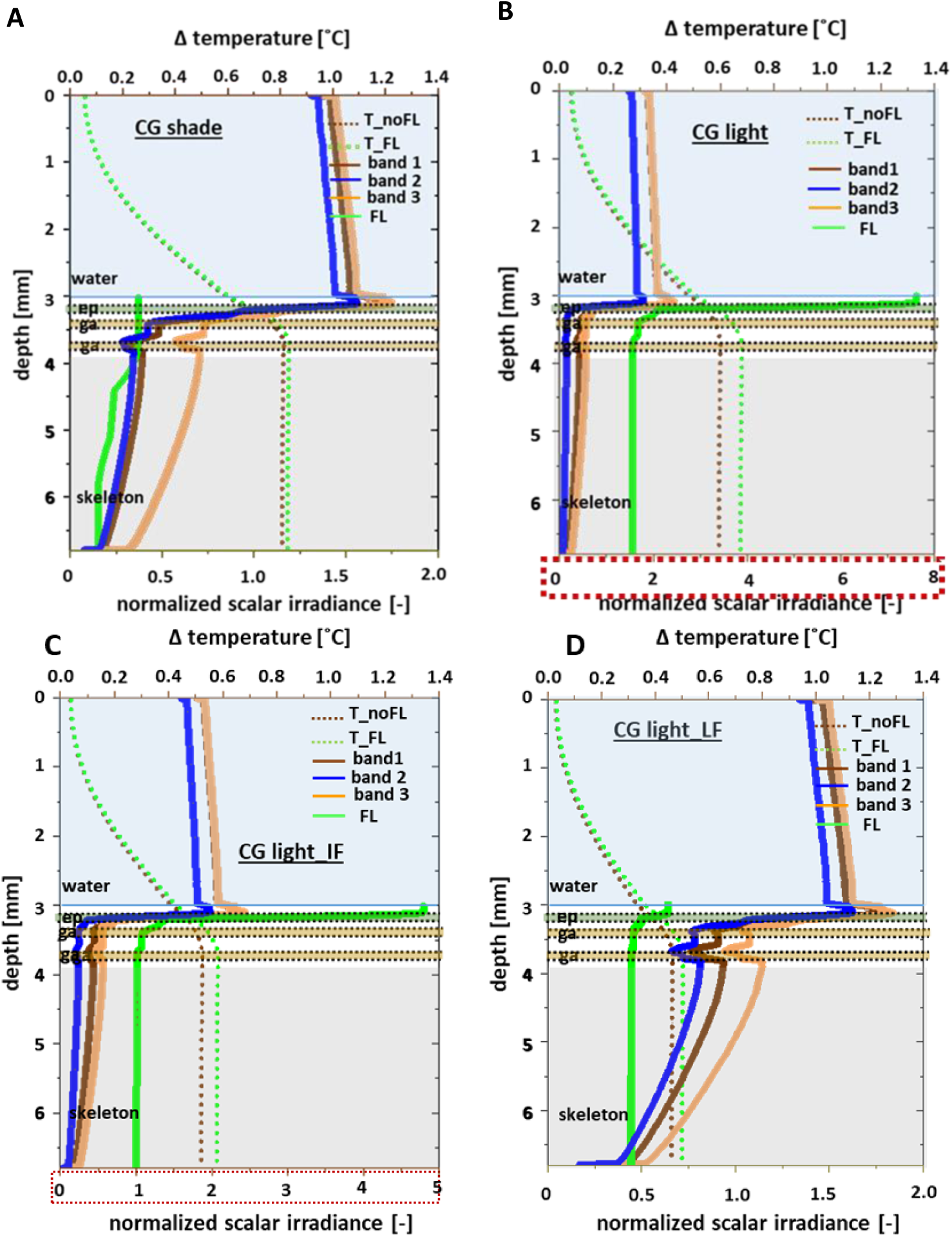
Optical and thermal effects of fluorescent granular protein (FP) expression and symbiont density in Cyphastrea microphthalma under contrasting light conditions and different FP concentrations. (A–B) Simulated depth profiles of normalized scalar irradiance (waveband-resolved) and temperature for CG (fluorescent) morphs under shaded and light-exposed conditions, respectively. Fluorescence emission (FL) was included as a secondary light source, and temperature profiles are presented both with (T_FL) and without (T_noFL) fluorescence re-emission.). (C) Simulated depth profiles of normalized scalar irradiance (bands 1–3 and fluorescence emission, FL) and temperature for CG_light fragment assuming a lower FP (IF) concentration, intermediate between that assumed for CG_shade and CG_light conditions shown in Fig. 4 (D) Simulated depth profiles of normalized scalar irradiance (bands 1–3 and fluorescence emission, FL) and temperature for CG_light assuming low FP concentration (LF, CG_shade concentration), while keeping the symbiont density identical to that of the measured CG light condition. Band 1: 400– 450 nm and 650–680 nm, principal absorption maxima of Symbiodiniaceae pigments; Band 2: 450–500 nm, absorption range for FP and Symbiodiniaceae pigments; band 3: rest; ep: epidermis; ga: gastrodermis. The directionality of scattering assumed here is 0.34.

**Figure S3.**
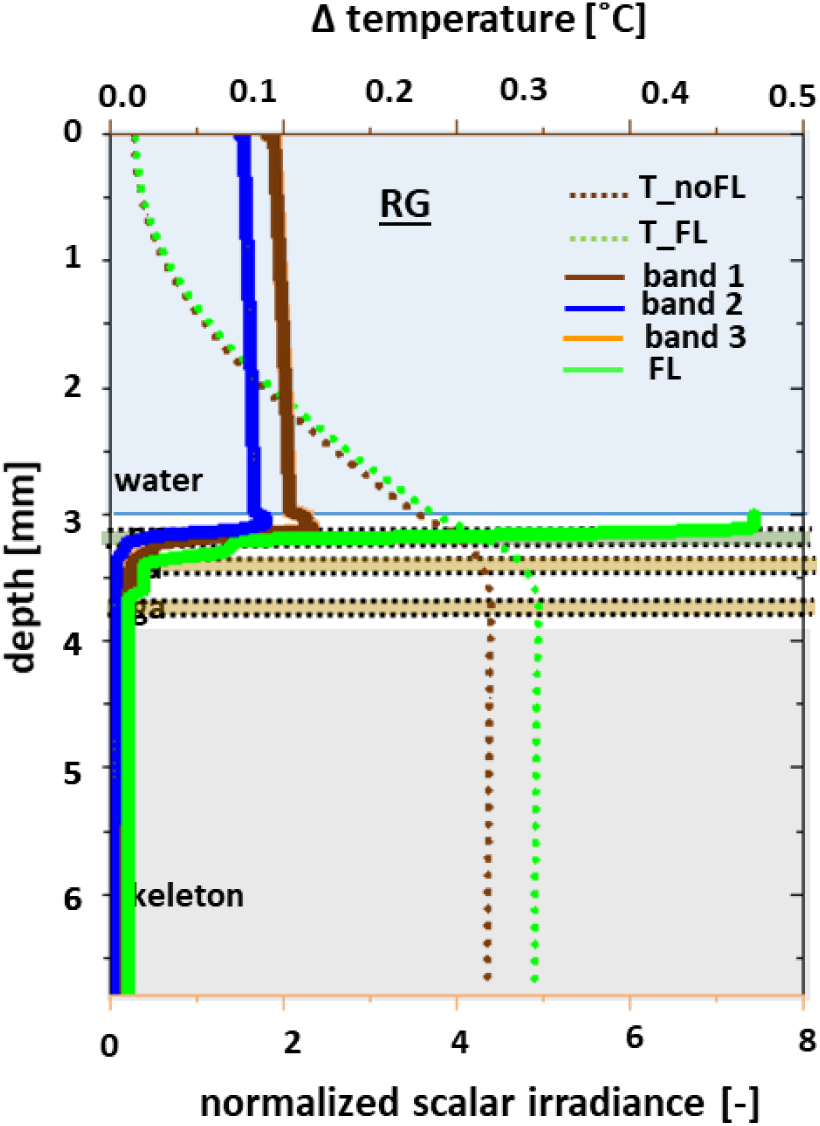
Optical and thermal effects of granular fluorescent proteins in A. curvata fragment. Simulated depth profiles of scalar irradiance (bands 1–3) and temperature for RG. Band 1: 400–450 nm and 650–680 nm, principal absorption maxima of Symbiodiniaceae pigments; Band 2: 450–500 nm, absorption range for FP and Symbiodiniaceae pigments; band 3: rest; ep: epidermis; ga: gastrodermis. g = 0.34.

**Figure S4:**
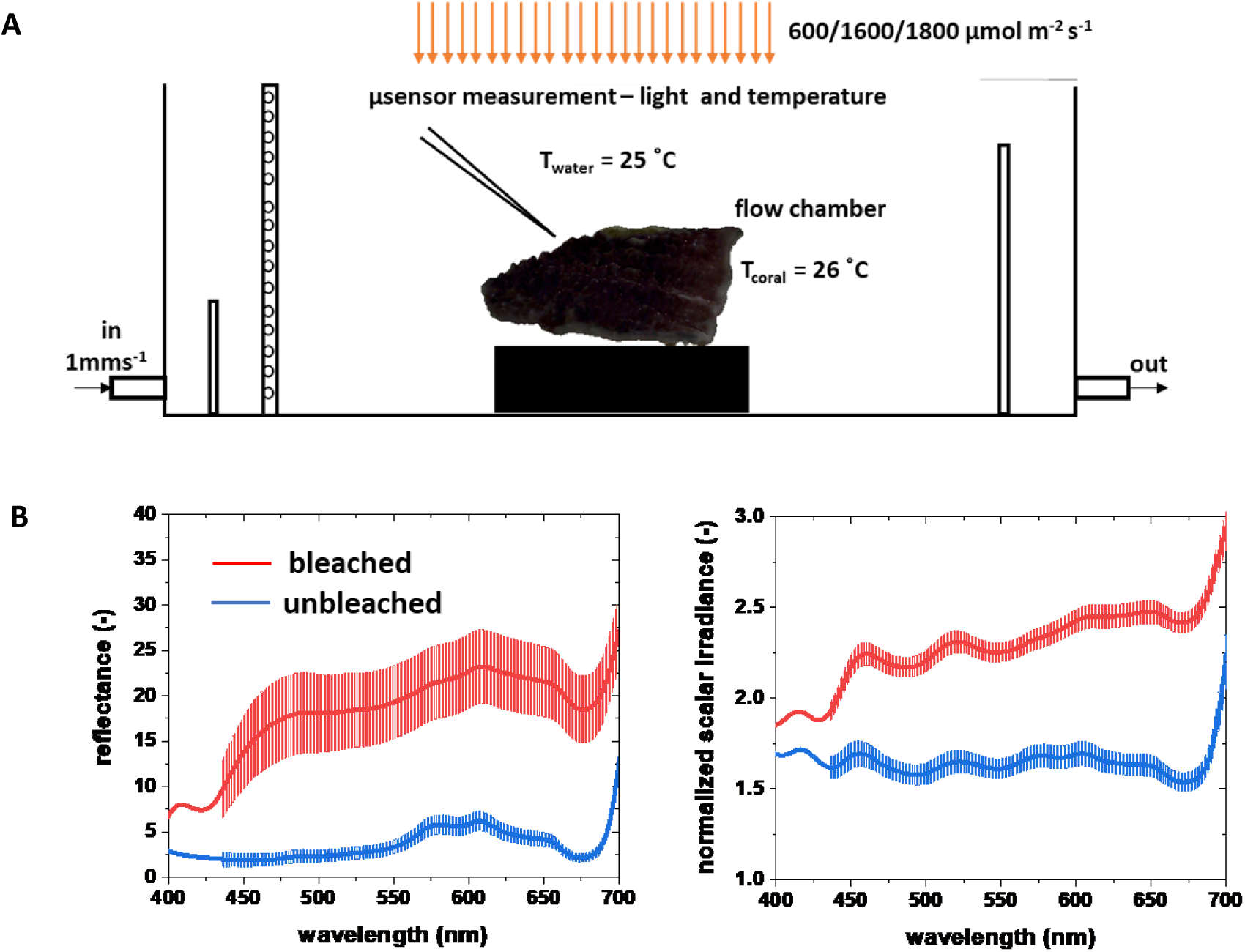
Overview of the experimental set-up. (A) Schematic of experimental flow chamber used for microsensor measurements of light and temperature at the surface of a coral fragment. Flow velocity was maintained at 1 mm s⁻¹. Fragments were exposed to different irradiance levels according to their prior acclimation history (field-collected or aquarium-reared). (B) Representative spectra of reflectance and scalar irradiance within the PAR range, measured using microsensors coupled to a spectrometer and normalized to the incident irradiance.

**Figure S5:**
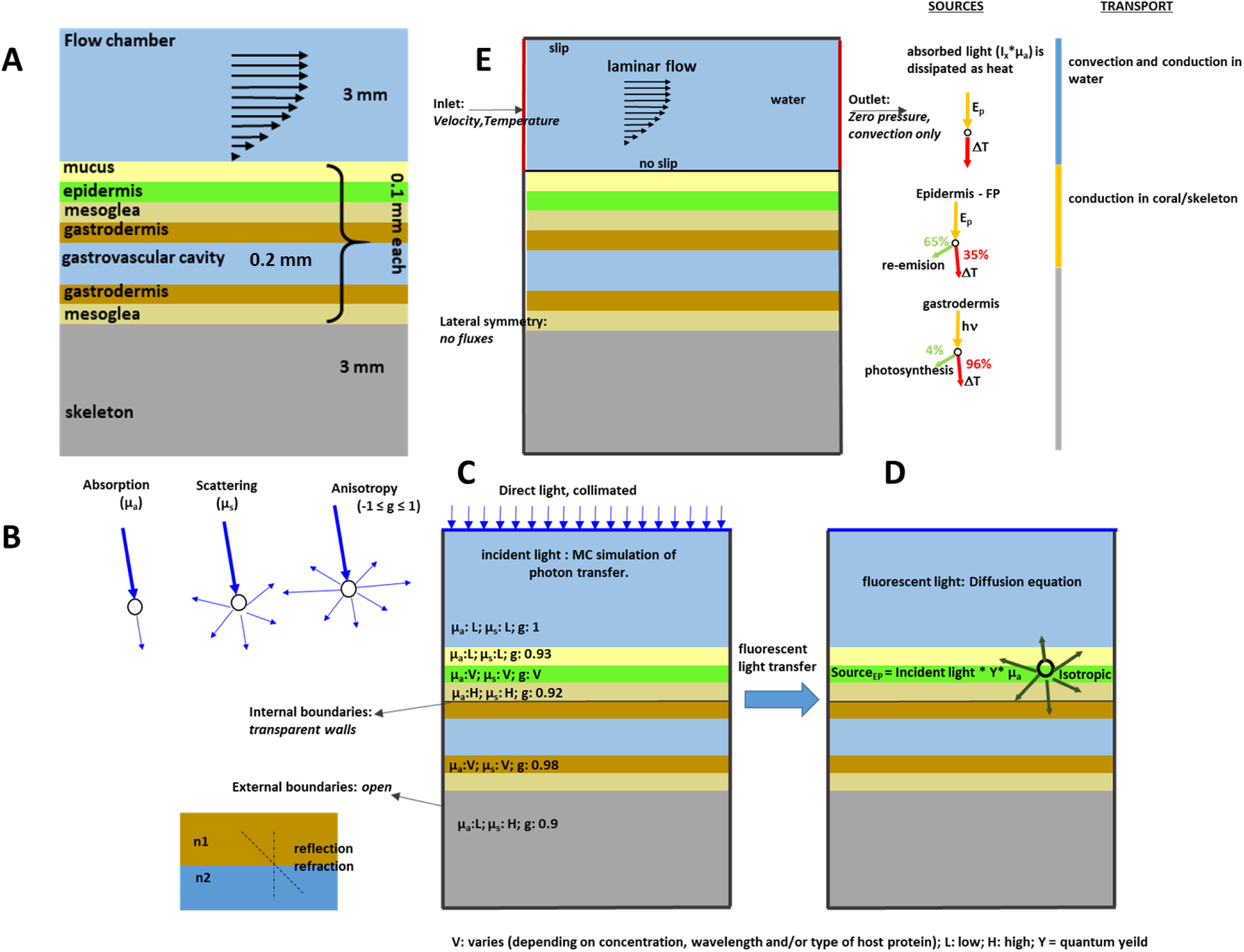
Overview of the simulation steps (A) 2D Coral slab model used or the simulations. The model represents the basic tissue organization of a coral, comprising a thin, stratified tissue layer overlying the skeleton. The geometry includes external medium (seawater), upper tissue layer, gastrovascular cavity, lower tissue layer, and coral skeleton. All sub-tissue layers are modeled with a thickness of 0.1 mm, except the gastrovascular cavity, which is set to 0.2 mm. The seawater and skeleton compartments are each modeled with a thickness of 3 mm. (B) Schematic illustrating the optical processes incorporated in the Monte Carlo simulation of light propagation, including absorption, scattering, scattering anisotropy, and Fresnel reflections at interfaces between layers with mismatched refractive indices (n). (C) Schematic illustrating the assumed optical properties under broadband (PAR) incident light, including the absorption coefficient (µ_a_), scattering coefficient (µ_s_), scattering anisotropy (g), and refractive index (n). The exact numerical values of these parameters are provided in Table X. In the schematic, the optical properties are shown qualitatively for clarity: L indicates low, H indicates high, and V indicates values that vary depending on the wavelength of the incident light and/or the physiological state of the coral fragment. (D) If the coral fragment contains a significant concentration of fluorescent proteins in the epidermis, a fraction of the incident light is re-emitted as fluorescence. This process is simulated by solving the diffusion equation, where the epidermal layer is defined as an isotropic light source. The source intensity is calculated as the product of the simulated incident light intensity (obtained from the Monte Carlo simulation), the fluorescence yield (Y), and the epidermal absorption coefficient (µ_a, epidermis_).

## Notes

### Competing Interest Statement

The authors have declared no competing interest.

